# Utilizing Genomics to Identify Novel Immunotherapeutic Targets in Multiple Myeloma High-Risk Subgroups

**DOI:** 10.1101/2025.01.29.635544

**Authors:** Enze Liu, Oumaima Jaouadi, Riya Sharma, Nathan Becker, Travis S. Johnson, Parvathi Sudha, Vivek S. Chopra, Faiza Zafar, Habib Hamidi, Charlotte Pawlyn, Attaya Suvannasankha, Rafat Abonour, Brian A. Walker

**Affiliations:** Melvin and Bren Simon Comprehensive Cancer Center, Division of Hematology and Oncology, School of Medicine, Indiana University, Indianapolis, IN, USA; Department of Biostatistics & Health Data Sciences, School of Medicine, Indiana University, Indianapolis, IN, USA; Genentech Inc., South San Francisco, CA, USA; Institute of Cancer Research, 15 Cotswold Road, London, UK

## Abstract

Immunotherapy is now standard of care for multiple myeloma, where the most common targets are B cell maturation antigen, CD38, and G protein-coupled receptor class C group 5 member D but strategies to identify additional targets are needed. We have utilized two large datasets of genomic data and integrated them with existing databases to identify expressed cell surface targets in myeloma patients. Importantly, we also identify targets specific to genomic defined subgroups of patients including primary translocations and high-risk subgroups. Examples of subgroup targets include *ROBO3* in t(4;14), *CD109* in t(14;16), *CD20* in t(11;14), *GPRC5D* in 1q+, and *ADAM28* in biallelic *TP53* samples. Expression was validated by flow cytometry and CRISPR-Cas9 knock out models. Sub-clonal differences in expression were noted, as was alternative splicing of existing immunotherapy targets such as *FCRL5*. These results highlight the use of genomic stratification to identify novel therapeutic targets.

**Graphical Abstract:** 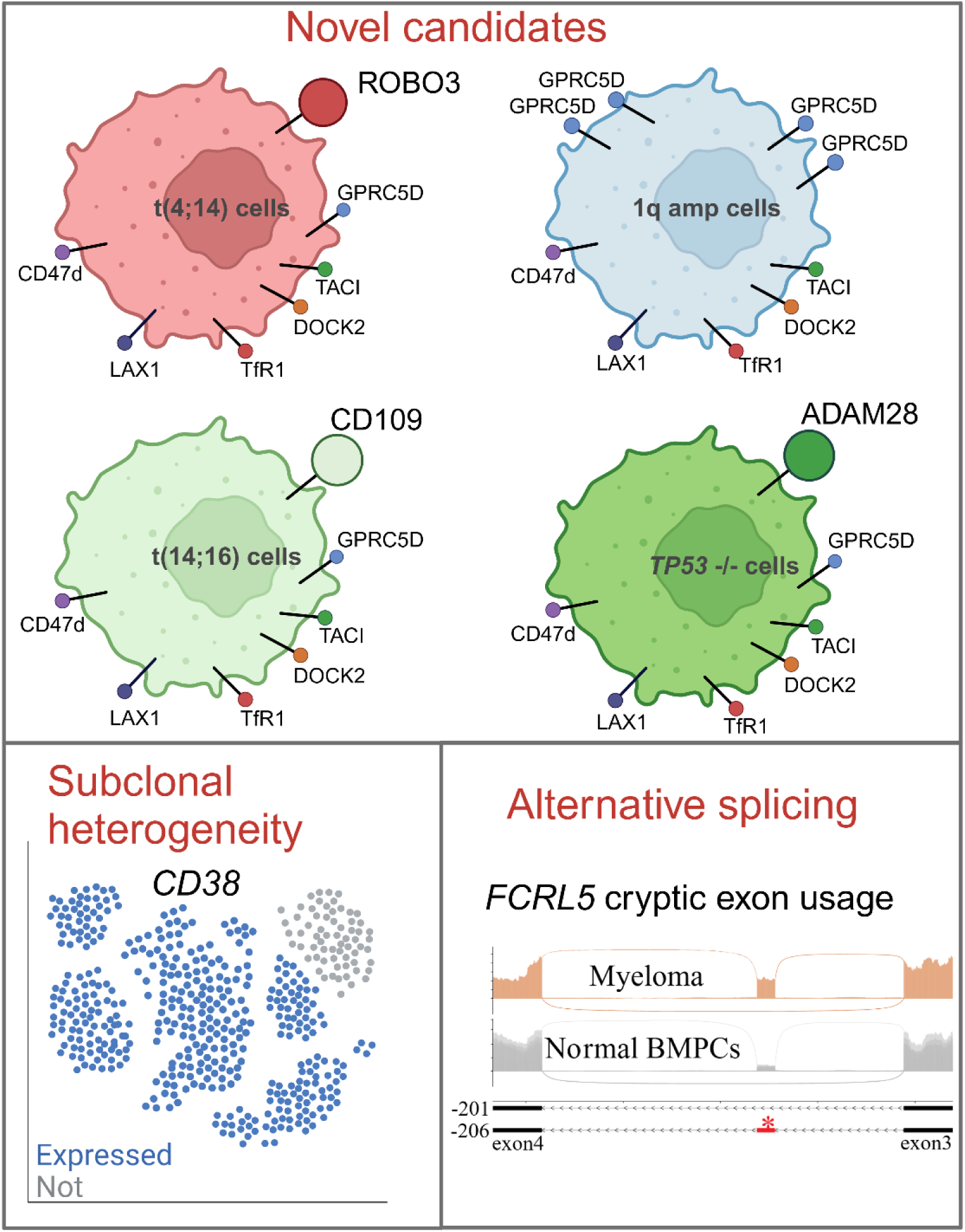

## Introduction

Multiple myeloma (MM) is the second most common hematologic malignancy. Despite advances in the treatment of patients, MM remains incurable due to the emergence of therapy resistance and disease relapse (1), indicating an unmet need for novel therapeutic options. Targeted immunotherapies have shown promise in killing MM cells with reduced toxicity towards normal cells (2), and therapies such as monoclonal antibodies (mAbs), antibody-drug conjugates (ADCs) and chimeric antigen receptor (CAR) T-cells/Natural killer (NK) cells have demonstrated efficacy and greatly prolonged patient survival (2–4).

The current cell surface targets on MM cells include B cell maturation antigen (BCMA), CD38, G protein-coupled receptor class C group 5 member D (GPRC5D), and Fc receptor-homolog 5 (FcRH5). Whilst targeting these proteins has proved to be successful, relapses still occur due to tumor intrinsic or extrinsic mechanisms. Tumor intrinsic mechanisms of resistance include deletion or mutation of the gene (5, 6), but could also include RNA splicing which has been seen in other B cell malignancies (7), Therefore, new targets are required to combat new clones that lose cell surface expression of the original target. Newer pan-MM targets identified include SEMA4A and ILT3 (8, 9), which give a wider repertoire of proteins to target multiple relapse patients.

Nevertheless, given that MM is a complex genomic disease there may be additional targets to be discovered. Firstly, MM is characterized by various subtypes, defined by five primary driver events and multiple secondary driver events (10). Different combinations of these genomic abnormalities may drive distinct surfaceome expression landscapes (11, 12). Secondly, heterogeneity has shown that subclones from the same patient exhibited different expression levels of these targets and could expand due to treatment selection (13). Additionally, identifying markers of high-risk disease could improve precision medicine whilst targeting the proliferative clone.

Here we use two datasets with genomic and transcriptomic data to systematically identify novel candidate targets with high potential to be used in immunotherapies. We designed a framework that utilized several knowledge bases from established proteomics experiments to select genes with high surface potential, which were further evaluated for toxicity in healthy organs and blood cells at both the population and subtype level. Population-based and subtype-based candidates were identified and validated by flow-cytometry, Western blot, and CRISPR-Cas9 knockout experiments. Additionally, single-cell data were utilized to determine their expression among subclones. Taken together, we identified novel cell surface candidates and validated several by experimental methods.

## Results

### Identification and characterization of surface targets

In order to identify potential immunotherapy targets against MM, we used a large dataset of RNA-sequencing from CD138+ sorted patient samples comprising of 837 newly diagnosed (ND) and relapsed/refractory (RR) samples from the MMRF CoMMpass dataset (MMRF) and 94 samples from the Indiana Myeloma Registry (IU dataset), plus eight normal bone marrow plasma cell samples (normal dataset) (**Table 1**). After data pre-processing, 19,892 protein-coding genes were annotated using five databases for cell surface potential. Those genes that encode proteins annotated with cell surface potential in ≥ three of those databases were taken further (n=845). Another 88 protein coding genes were added from proteomic studies where they were detected on the surface of MM cells (14), resulting in a total of 933 genes (**Figure 1A**).

**Figure 1.**
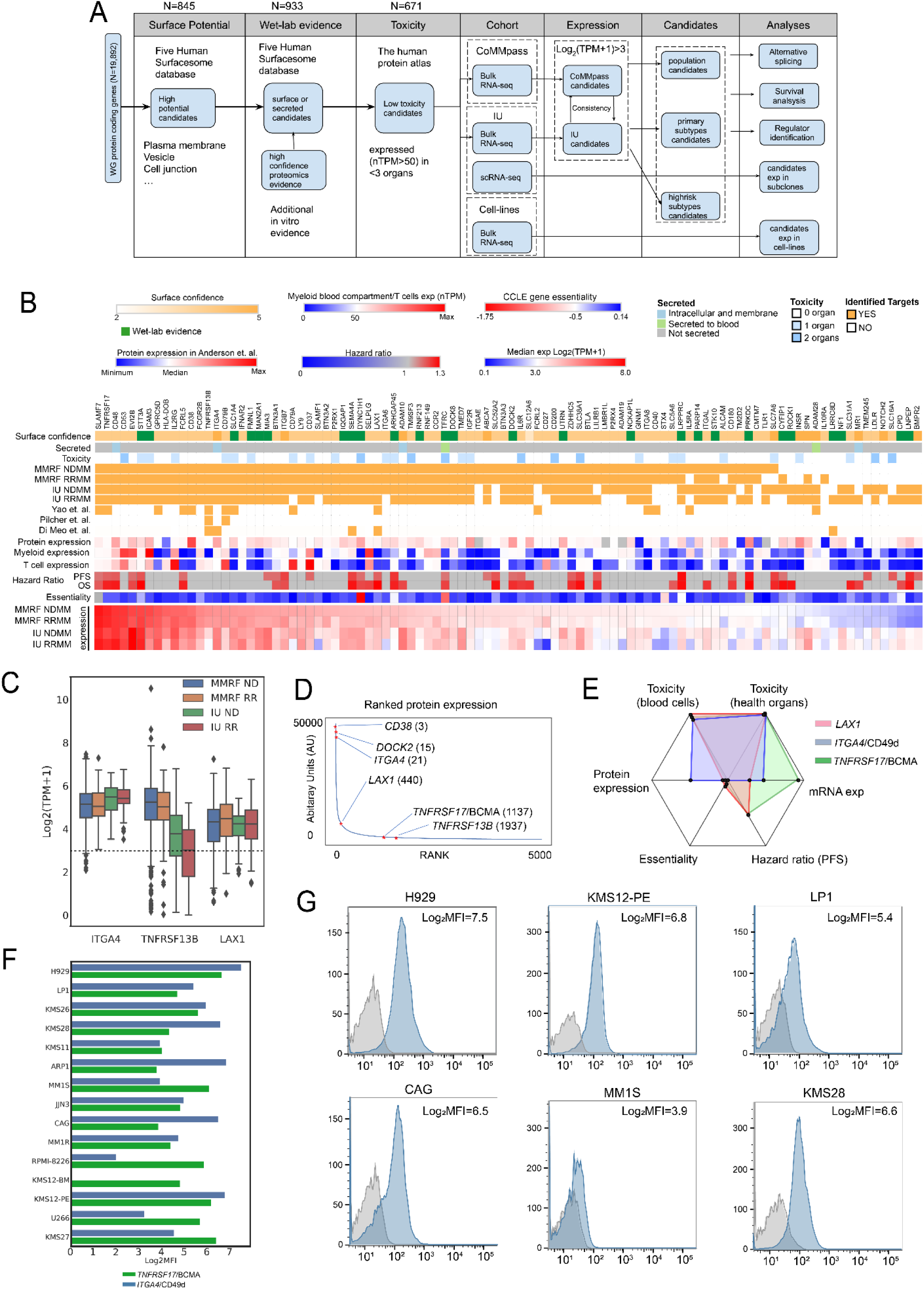
Characteristics of candidate targets identified in ND and RR populations from two independent datasets. (A) General workflow of the target identification process. (B) A heatmap demonstrating all identified candidates in ND and RR population from MMRF and IU datasets with various annotations. (C) Expression level of selected candidates. (D) Ranked expression of 5,092 proteins documented in Anderson et. al(8). Numbers after gene names: rank. (E) A radar plot summarizing key characteristics among *LAX1*, *ITGA4* and *TNFRSF17*/BCMA. Range (from center to edge): toxicity (healthy organs): 2∼0; toxicity (blood cells): 0∼1845; protein exp: 24.8∼37221.6; essentiality: 0∼ −1.75; hazard ratio (PFS): 1∼1.28; mRNA exp: 3∼7.6. Range in toxicity, protein expression, essentiality, hazard ratio and mRNA exp indicated lowest to highest among 98 candidates. (F) Log_2_-scaled median fluorescence intensity (MFI) of TNFRSF17/BCMA and ITGA4/CD49d detected by flow cytometry in 15 MM cell lines. (G) Density plots indicating MFI (blue peaks) of *ITGA4*/CD49d compared to the isotype control (grey peaks) across 6 MM cell lines. Log_2_MFI: Log_2_-scaled MFI.

**Table 1.** Dataset summary.

|  |  | MMRF |  |  | IU |  |  |
| --- | --- | --- | --- | --- | --- | --- | --- |
|  |  | NDMM | RRMM | Total | NDMM | RRMM | Total |
| Primary subtypes | **Total** | **754** | **83** | **837** | **44** | **50** | **94** |
|  | **Translocations** |  |  |  |  |  |  |
|  | t(14;16) | 31 | 2 | 33 | 0 | 3 | 3 |
|  | t(14;20) | 10 | 2 | 12 | 0 | 2 | 2 |
|  | t(4;14) | 92 | 4 | 96 | 3 | 12 | 15 |
|  | t(11;14) | 134 | 14 | 148 | 14 | 12 | 26 |
|  | non-translocated | 471 | 61 | 532 | 26 | 20 | 46 |
|  | other | 16 | 0 | 16 | 1 | 1 | 2 |
|  | **Hyperdiploid Status** |  |  |  |  |  |  |
|  | Hyperdiploid | 384 | 36 | 420 | 23 | 20 | 43 |
|  | Non-Hyperdiploid | 276 | 28 | 304 | 21 | 30 | 51 |
|  | Undetermined | 94 | 19 | 113 | 0 | 0 | 0 |
| High-risk subtypes | **TP53 status** |  |  |  |  |  |  |
|  | biallelic *TP53* | 23 | 4 | 27 | 4 | 11 | 15 |
|  | mono allelic *TP53* | 62 | 11 | 73 | 3 | 5 | 8 |
|  | *TP53* WT | 549 | 42 | 591 | 37 | 34 | 71 |
|  | undetermined | 120 | 26 | 146 | 0 | 0 | 0 |
|  | **1q status** |  |  |  |  |  |  |
|  | 1q amp | 39 | 5 | 44 | 7 | 13 | 20 |
|  | 1q gain | 182 | 20 | 202 | 13 | 14 | 27 |
|  | 1q WT | 413 | 31 | 444 | 24 | 23 | 47 |
|  | undetermined | 120 | 27 | 147 | 0 | 0 | 0 |
|  | **GEP group** |  |  |  |  |  |  |
|  | PR | 24 | 13 | 37 | 3 | 10 | 13 |
|  | MF | 35 | 4 | 39 | 0 | 4 | 4 |
|  | MS | 96 | 7 | 103 | 5 | 6 | 11 |
|  | Other subgroups | 599 | 59 | 658 | 36 | 30 | 66 |

To reduce potential toxicities from off-target effects, these 933 genes were further filtered based on their expression in normal organs from the Human Protein Atlas. After examining the expression of several existing CAR-T/immunotherapy targets in MM and other blood cancers (**Supplementary Figure 1**), we established a threshold of nTPM>50 in no more than two essential organs. Such criterion effectively encompassed all known targets while maintaining specificity to reduce off-target risks. The last filter applied was based on gene expression in patient MM cells (log_2_(TPM+1)>3). This threshold was integrated into the selection process, which resulted in 98 candidate genes (**Supplementary Figure 1**).

Of those 98 candidates, 74 were expressed at log_2_(TPM+1)>3 in both datasets (**Figure 1B**) and 88 (91.7%) were consistently expressed in both newly diagnosed and relapsed samples. Approved immunotherapy targets ranked highly by their RNA expression level including *SLAMF7* (rank 1 in MMRF newly diagnosed MM), *TNFRSF17*/BCMA (rank 2), *GPRC5D* (rank 8), *FCRL5* (rank 11), and *CD38* (rank 12), and were identified in all datasets and disease stages. Recently reported novel candidates(14) such as *CD48*(*15*) (rank 3), *CD53*(*16*) (rank 4), *EVI2B*/CD361(16) (rank 5), *CD79A/B*(*17*) (rank 16 & 24), *SEMA4A*(*8*) (rank 31), and *LILRB4*/ILT3(9) (rank 60) were also identified using this methodology. Other previously identified candidates such as *STT3A*(*18*) (rank 6) and *ICAM3*(*16*) (rank 7) were not annotated as membrane proteins in public databases but were rescued by our methodology as they had been detected as cell surface proteins in proteomic experiments (8). An additional level of complexity is reflected in the differences between RNA and protein expression levels (r<0.3, Spearman correlation) (**Supplementary Table 2**), showing the need for matched RNA and protein level data in the same samples to utilize existing genomic data more effectively.

Clinical evidence suggests that maintaining a basic level of immunity may reduce infection-induced mortality in myeloma patients (19, 20). To this end, we further examined the expression of targets in myeloid cells and found that their expression level varied (**Figure 1B** and **Supplementary Figure 1**). *GPRC5D* and *FCRL5* had minimal expression in myeloid cells, *SLAMF7*/CD319 and *TNFRSF17*/BCMA had moderate expression, whereas *CD53*, *CD48* and *EVI2B* exhibited highest expression.

40 candidates were found to have significant associations with either progression-free survival (PFS) or overall survival (OS) in the MMRF newly diagnosed dataset (**Figure 1B**), including 25 candidates that were associated with both (**Supplementary Table 2**). This included established targets such as *TNFRSF17*/BCMA as well as novel targets such as *LAX1, ITGA4, DOCK2, and TNFRSF13B/*TACI (**Figure 1C-D, Supplementary Figure 2**). CD27 has also been targeted together with CD70 in bispecific therapies in solid tumors and AML (21–23).

*LAX1* encodes a lymphocyte transmembrane adaptor protein involved in B cell activation, which had high mRNA (**Figure 1C**) and protein expression (440^th^, **Figure 1D**) across samples. Its expression showed significant association with both PFS and OS (Hazard ratio (HR)=1.2 and 1.3, p=0.01 and p<0.005, **Figure 1B & Supplementary Figure 2**). *ITGA4* encodes a subunit of integrin alpha 4 chain found at the surface of immune cells and plays a crucial role in cell adhesion and migration. Several FDA-approved immunotherapies target it in autoimmune diseases such as ulcerative colitis (24), showing its accessibility as a target. Compared to BCMA, *ITGA4* had higher expression at the proteomic level, low toxicity and low off-target effect (**Figure 1E**). Moreover, flow cytometry experiments suggested that *ITGA4*/CD49d was accessible and highly expressed across most of the MM cell lines, even compared to TNFRSF17/BCMA (**Figure 1F-G**).

### Identifying targets essential for MM cell survival

Despite the effectiveness of immunotherapy in treating MM patients, eventually tumor cells can mutate or delete the epitopes in the targets, resulting in relapse. Current immunotherapy targets are not essential for MM cell growth, but it may be possible to identify targets that are essential for growth, thereby limiting tumor intrinsic resistance mechanisms. Six of the identified candidates (*DYNC1H1, ARHGAP45, TFRC, STX4, LRPPRC* and *CPD*) demonstrated significant essentiality (median CERES(25) gene effect score<-0.5) in 18 MM cell lines, of which three (*DYNC1H1, TFRC* and *LRPPRC*) were significantly associated with patient survival (**Figure 1B**). Many of the encoded proteins are likely to only be associated with the cell membrane, or not have domains that extend into the extracellular space. The exception was *TFRC*, the transferrin receptor, which had high essentiality (median CERES<-1), is involved with iron intake, its expression was previously reported to be associated with disease progression in MM (26), and it has been reported as a potential target in hematological malignancies (27). *TFRC* expression was significantly associated with patient survival (**Supplementary Figure 3**), was expressed (nTPM>50) in one off-target organ (ovary) and was not expressed in myeloid cells (**Supplementary Figure 3**) suggesting its potential as a candidate with low side-effects. However, others have reported that *TFRC* expression is not specific to plasma cells (28). Our findings suggest that, besides their oncogenic roles in MM cells, these essential genes encoding cell surface proteins could be targeted by immunotherapies, but largely cell surface proteins are not essential for survival of MM cells.

### Heterogeneity of immunotherapy target expression in MM subgroups

Given that we know MM is a heterogeneous disease driven by primary genomic abnormalities, we utilized expression of the 98 candidates as features and conducted clustering analysis for all samples in the two datasets (**Figure 2A**). A clear segregation among t(11;14), t(4;14), t(14;16), t(14;20) and non-translocated subgroups was observed, indicating heterogeneity of candidate target gene expression. However, no clear segregation was observed between ND and RR timepoints, and so these were merged in subsequent analyses.

**Figure 2.**
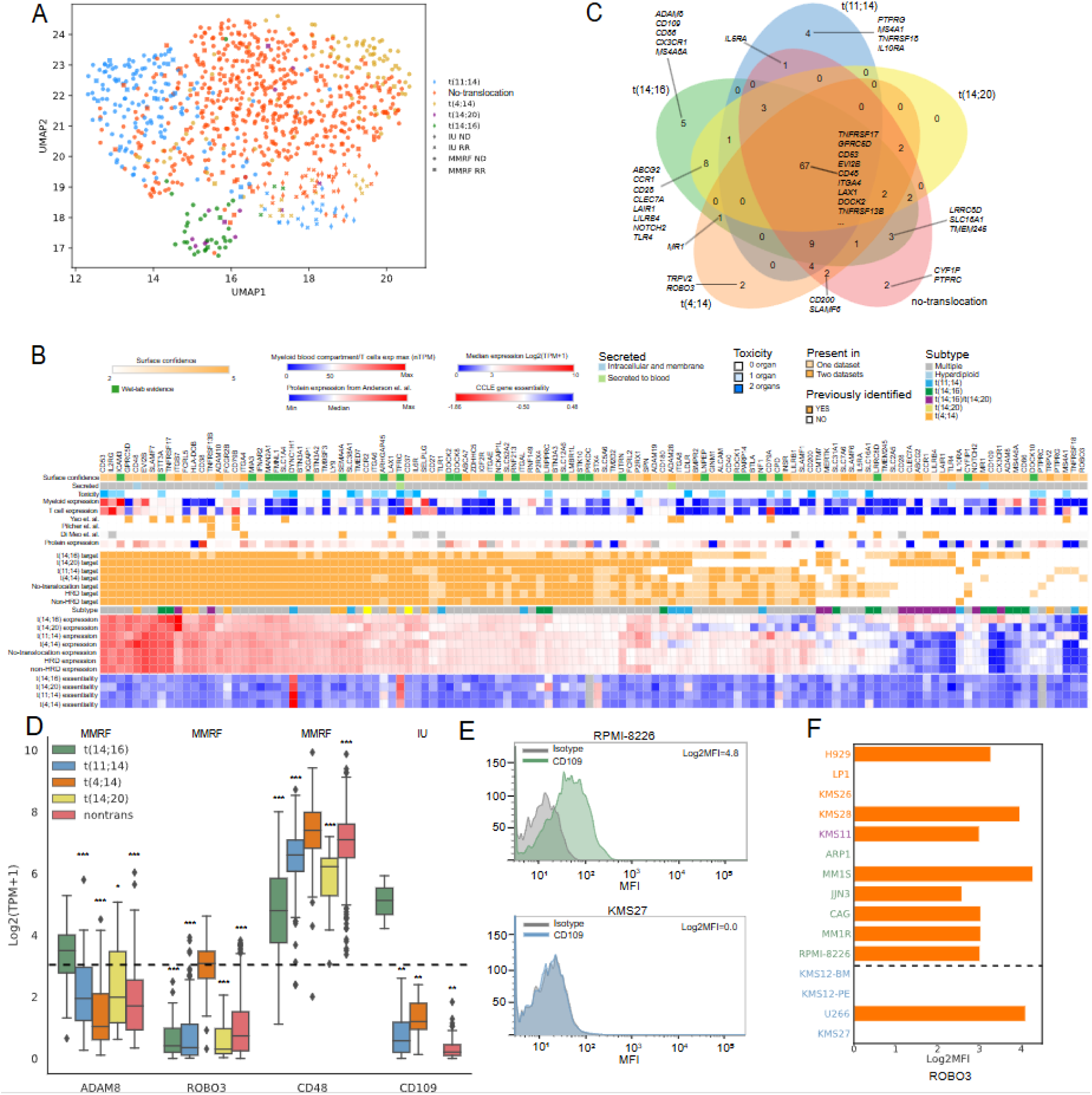
Characteristics of candidates identified from primary subtypes. (A) UMAP plot indicating the expression heterogeneity of 98 population-based candidate targets among five primary subtypes. (B) A heatmap demonstrating all identified candidates from primary subtypes. (C) A Venn diagram indicating the common/unique candidate targets among primary subtypes. (D) Expression level of selected candidate targets uniquely/highly expressed in certain subtypes. (E) Protein expression of *CD109* was detected in RPMI-8226 (t(14;16)) but not in KMS27 (t(11;14)) cell line. (F). Protein expression of *ROBO3* and *TNFRSF17*/BCMA in t(4;14) (orange), t(14;16) (green), and t(11;14) (blue) cell lines. Statistical tests in (D): Mann-Whitney U test; Significance level: * p<0.05; ** p<0.01, *** p<0.001.

Samples were annotated with their primary genomic events and the same target selection process was performed on each subgroup, including sub-group specific expression, dependency, and outcome (**Figure 1A**). The subgroup analysis identified 120 candidate targets (**Figure 2B-C, Supplementary Figure 4 and Supplementary Table 3**). Among them, 22 candidates were only identified in the subgroup analysis, indicating the advantage of using a subgroup approach. Over half (67/120) of the candidate target genes were expressed across all subgroups and included the established pan-MM candidates such as *TNFRSF17*/BCMA, *SLAMF7*, *GPRC5D*, *CD38*, and *FCRL5*, as well as novel candidates such as *LAX1*, *ITGA4* and *TNFRSF13B (TACI)* (**Figure 2B-C**).

A total of 21 candidate targets were only expressed above the cut-off within a subtype. Most unique candidates were found in the t(MAF) (t(14;16) or t(14;20)) subgroups (N=13) followed by the t(11;14) subgroup (N=4) (**Figure 2B-C**). Although *ITGB7* was identified as a pan-myeloma candidate, as previously noted(26) it was expressed at much higher levels in the t(MAF) subgroups (**Figure 2B**), and has been identified as a CAR-T cell target (29). Similarly, *CD79A and CD20/MS4A1* had higher expression in the t(11;14) subgroup and have been used as immunotherapeutic targets in other hematological malignancies.(30, 31) Other subgroup-specific candidates have also been identified as immunotherapy targets in other diseases, including *LILRB4* (t(MAF)-specific) which was recently established as novel CAR-T target in AML.(32) *CLEC7A* (MAF-specific),(33) *CD86* (t(14;16)-specific) (34), and *CD28 (t(MAF)-specific)* (35) have been applied as CAR-T targets in solid tumors.

Heterogeneity of expression between patient samples and cell lines was also observed, even for established targets (**Supplementary Figure 4**). For instance, *FCRL5*, *EVI2B,* and *CD27* had significantly higher RNA expression (Log_2_FC=4.3, 3.9, 2.5, p=3×10^−12^, 2×10^−12^, 4×10^−8^, respectively) in patient samples than cell lines, while *CD70*, which was a partner of *CD27* in bispecific therapies (21–23), showed a significantly higher expression in cell lines (Log_2_FC=3.7, p=3×10^−11^). Meanwhile, candidates such as *ARHGAP45* and *TNFRSF17* showed no significant difference in expression, suggesting a more uniform expression of these candidates in different models.

Of the 120 candidate targets identified at the subgroup level, 23 demonstrated significant associations between expression and both PFS and OS (**Supplementary Table 3**). High expression of one (*LAIR1* in the MAF subgroup) was associated with a good prognosis and the others were associated with a poor prognosis. 19 candidate targets showed an association with outcome in only one subtype, indicating their subtype specificity. For example, *CD180* was significantly associated with inferior PFS and OS only in the hyperdiploid subtype (HR=1.2 and 1.3, p=0.01 and p=0.02, respectively, **Supplementary Figure 5**), indicating its potential specificity in maintaining hyperdiploid cell proliferation. This gene encodes a component of the cell surface receptor complex RP105/MD-1 (36), which is associated with hematological malignancies (37). It has low expression in normal tissues and blood cells (**Supplementary Figure 5**) suggesting its validity of a candidate.

We speculated that t(14;16)/t(14;20)-specific candidates were uniquely regulated by their master regulator *MAF*/*MAFB* and therefore analyzed existing ChIP-seq data of *MAF*/*MAFB* binding sites.(38) Four of the t(*MAF*)-specific candidates were found to have MAF binding sites (**Supplementary Figure 5**), including at the promoter region of *ITGB7* and *CD109* (**Supplementary Figure 5**). The former has been well-documented as over-expressed in the t(*MAF)* subgroup and as a target of *MAF* transcription families (39), while the latter was found to be expressed in solid tumors (40). Moreover, *CD109* encodes a glycoprotein on the surface of platelets (36) and was solely expressed in the t(14;16) subgroup (**Figure 2D**), and its expression in healthy organs and blood cells was minimal (**Supplementary Figure 5**). Flow cytometry showed expression of *CD109* in t(14;16) and t(4;14) cell lines (especially in RPMI-8226), but not in t(11;14) cell lines (Log_2_FC=3.8, p=0.03, one-sided Mann Whitney U test, **Figure 2E-F**, **Supplementary Figure 5**).

Other notable subtype-specific candidates included *ROBO3*. This gene was selected due to its specific expression in t(4;14) subgroups (Log_2_FC>2, p<1×10^−8^, MMRF dataset, **Figure 2D**) and its minimal expression in healthy organs and blood cells (**Supplementary Figure 5**). Flow cytometry validated the expression of *ROBO3* in several t(4;14) and t(14;16) cell lines and lack of expression in other cell lines (Log_2_FC=1.3, p=0.17, one-sided Mann Whitney U test, **Figure 2F**). Additionally, *CD48* had been identified as a common candidate across all cytogenetic groups but its expression was significantly higher in the t(4;14) subtype (Log_2_FC>0.3, p<0.0009, MMRF dataset, **Figure 2D**). Higher expression plus low toxicity (**Supplementary Figure 5**) makes *CD48* a valid candidate for further exploration.

### High-risk subtypes are associated with higher expression of specific targets, including GPRC5D

High-risk secondary events such as 1q gain/amplification and biallelic *TP53* inactivation have been highly associated with inferior survival (41), disease progression/relapse (41), and therapy resistance (42). To evaluate novel targets in these high-risk subgroups we identified candidates in samples with 1q gain or amplification, *TP53* abnormalities and established high-risk expression subgroups (PR/MF/MS) (26). In total, 125 candidates were identified (**Figure 3A & Supplementary Table 4**), most of which overlapped with the previously identified candidates.

**Figure 3.**
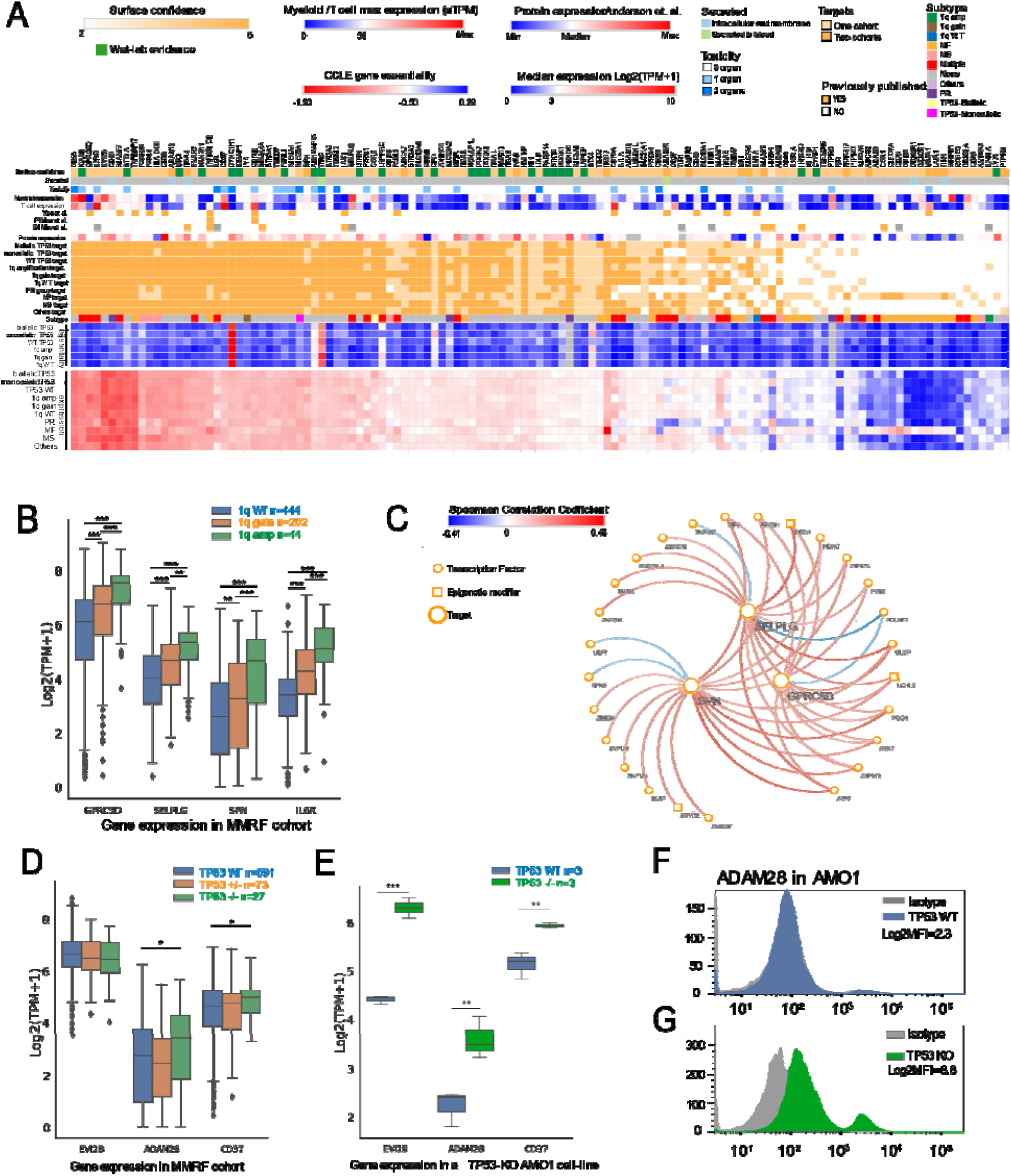
Characteristics of candidates identified from high-risk subtypes. (A) Identified candidate targets for high-risk subtypes. (B) Four candidates demonstrating significantly elevated expression levels along with 1q copy number gain in the MMRF dataset. (C) A network plot indicating potential regulators of *SELPLG*, *SPN* and *GPRC5D* found on 1q. (D) Three candidates demonstrating heterogenous expression among biallelic (−/−), monoallelic (+/−) and wild type (WT) *TP53* subgroups in the MMRF dataset. (E) Three candidates exhibiting heterogenous expression in *TP53* knock-out and WT triplicates of the AMO1 cell line. Protein expression of ADAM28 detected in TP53 wild-type (F) and knockout (G) AMO1 cell line measured by flow cytometry. Statistical test in boxplots: Mann-Whitney U test in (B) and (D),T-test in (E). significance level: * p<0.05, ** p<0.01, *** p<0.001.

Since high-risk patients have been shown to develop a PR expression signature over time, they are an interesting group to identify therapeutic targets (12, 43). Three unique candidates were found in the PR subgroup: *CD300A*, *F2R* and *PTPRG*. *CD300A* was identified in the MMRF dataset while the other two were identified in the IU dataset. The three candidates showed minimal expression in healthy organs and *PTPRG* and *F2R* also had minimal expression in all blood cells (**Supplementary Figure 6**).

Several candidate targets had higher expression in samples with gain/amp1q (**Figure 3B**) including *GPRC5D*, *SELPLG*, *SPN*, and *IL6R*. *GPRC5D*, an established target in CAR-T therapy against MM (44), showed increased expression along with 1q gain in both the MMRF (1q WT vs. 1q gain, Log_2_FC=0.7, p=7×10^−8^; 1q gain vs. 1q amp, Log_2_FC=0.8, p=1×10^−3^, **Figure 3B**) and the IU datasets (**Supplementary Figure 7**). *SELPLG* (1q WT vs. 1q gain, Log_2_FC=0.6, p=7×10^−8^; 1q gain vs. 1q amp, Log_2_FC=0.6, p=6×10^−4^) and *SPN* (1q WT vs. 1q gain, Log_2_FC=1.2, p=4×10^−3^; 1q gain vs. 1q amp, Log_2_FC=1.2, p=2×10^−5^) encode glycoproteins that can lead to activation of T cells (45). *IL6R* is located on 1q and encodes the receptor of interleukin 6, which is a growth factor of MM cells (46). There was a step-wise increase in expression of *IL6R* with copy number gain (1q WT vs. 1q gain, Log_2_FC=0.9, p=2×10^−19^; 1q gain vs. 1q amp, Log_2_FC=0.9, p=5×10^−6^) indicating a gene dosage effect.

The other three genes upregulated with gain/amp1q were not located on 1q thus their elevated expression might be due to potential regulators present on 1q. Hence, we conducted a transcription factor-target analysis to infer the potential transcription factor(s) or epigenetic modifier(s) that could control expression of these genes. Among 1,629 human transcription factors (36), 55 were located on 1q. Three transcription factors had a high correlation of expression with *GPRC5D*, *SELPLG*, and *SPN* which included *ARNT* (Spearman correlation ρ=0.21, 0.25 and 0.33, rank: 9^th^, 10^th^ and 4^th^, vs. *GPRC5D*, *SELPLG* and *SPN,* respectively), *ATF6* (ρ=0.25, 0.36 and 0.42, rank: 7^th^, 3^rd^ and 1^st^) and *GLMP* (ρ=0.32, 0.45 and 0.25, rank: 1^st^, 1^st^ and 12^th^) (**Figure 3C, Supplementary Figure 7**). *ARNT* and *ATF6* had documented binding sites on *SELPLG* and *SPN* (36). Additionally, *PBX1* was recently found to be associated with 1q gains and tumor progression (47), although its correlation was not among the highest (p=0.15, 0.20 and 0.24, rank: 18^th^, 13^th^ and 13^th^, **Supplementary Figure 7**).

Biallelic *TP53* is arguably the most important genomic risk factor in MM (41). Comparisons of wild type *TP53* samples with biallelic *TP53* abnormalities identified 12 candidates with higher expression. For instance, *ADAM28* (Log_2_FC=0.5, p=0.04, one-sided Mann-Whitney U test) and *CD37* (Log_2_FC=0.4, p=0.05) showed higher expression in biallelic *TP53* vs. wild type patients in the MMRF dataset (**Figure 3D**).

Using existing expression data from isogenic AMO1 MM cells, where *TP53* had been knocked out (KO) by CRISPR/Cas9, *ADAM28* and *CD37* were expressed at significantly higher levels in *TP53* KO compared to WT cells (Log_2_FC=1.4 and 0.8, p=0.006 and p=0.004, one-sided T-test; **Figure 3E**). To determine if this effect was recapitulated at the protein level, we generated a *TP53* KO using the AMO1 cell line (absence of p53 confirmed by Western blotting, **Supplementary Figure 7**) and measured protein expression by flow cytometry. We validated *ADAM28* and confirmed its increased expression on the cell surface of *TP53* KO AMO1 cells compared to WT AMO1 (Log_2_MFI=6.0, Log_2_FC=3.7) (**Figure 3F-G**). ADAM28 is a member of the metalloproteinase-type A disintegrin and metalloproteinases (ADAMs) family and previously found to be upregulated in various cancers to promote cell proliferation, migration and invasion (48). This is a prime example of how integration of patient data with experimental models can be used to identify and validate potential future immunotherapy targets, especially in high-risk disease.

### Heterogeneous expression of candidate targets among subclones

To explore the expression of targets across all cells within a patient, we utilized scRNA-seq data to examine subclones within samples (**Figure 4A, Supplementary Figure 8**). Single-cell data was clustered allowing the expression of targets to be measured as a proportion of each patient. Several population level candidates were ubiquitously expressed including *TNFRSF17*/BCMA (94% of patients) and *SLAMF7* (93%, **Supplementary Figure 8**) as well as novel candidates including *ITGA4* (100%). Other candidates such as *CD79A* were preferentially expressed in more B-cell like subtypes including t(11;14) and hyperdiploid samples with 11q gain (26%, p=0.04, hypergeometric test, **Figure 4B**), reflecting inter-patient and subtype-specific expression heterogeneity of identified candidates.

**Figure 4.**
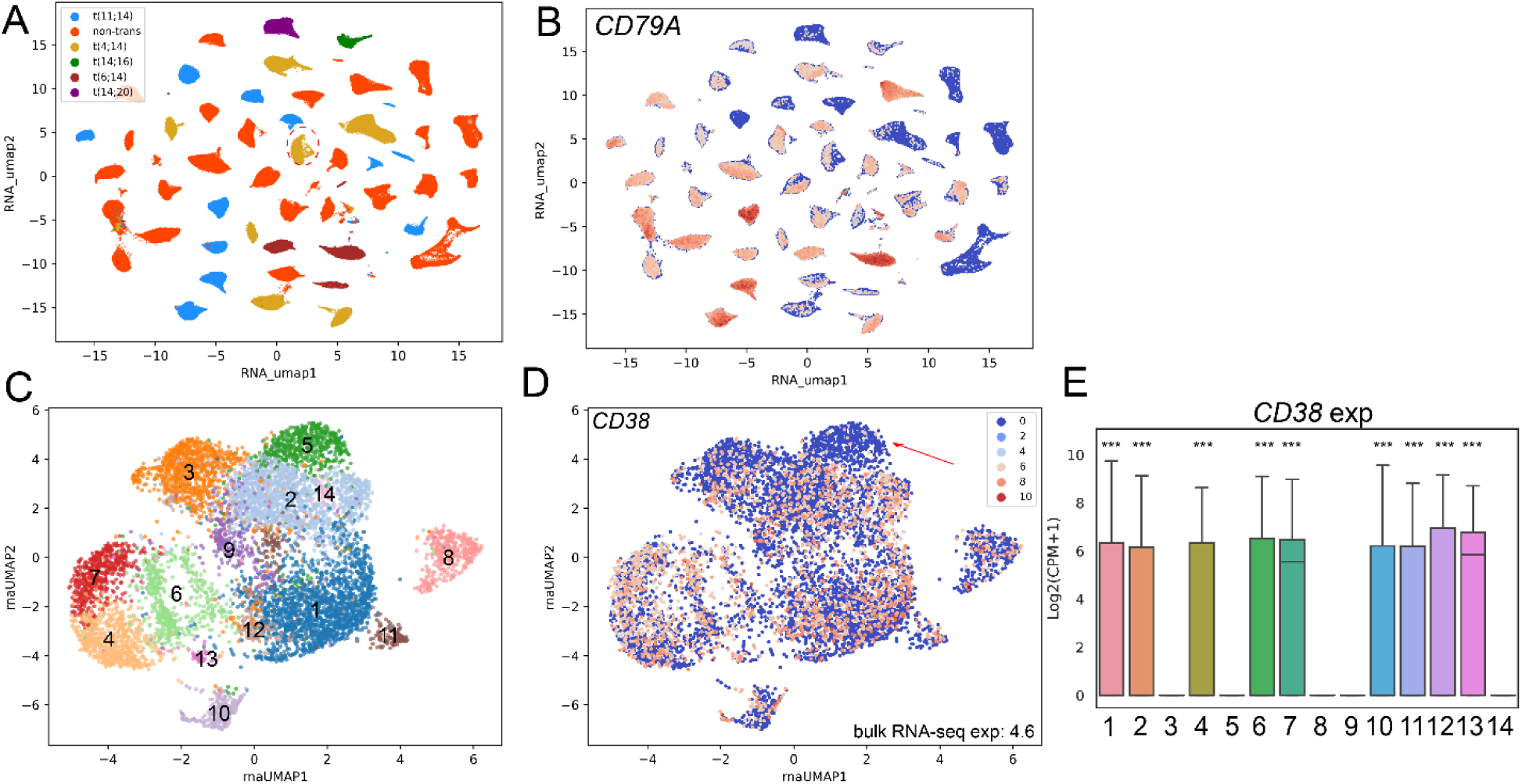
Candidate targets were heterogeneously expressed among subclones identified from scRNA-seq data. (A) Transcriptomic landscapes of 49 IU patient samples with single-cell RNA-seq data. (B) Expression of *CD79A* in is increased in t(11;14) and hyperdiploid 11q+ samples. (C) 14 subclones identified by scRNA-seq in a relapsed t(4;14) sample. (D) Uneven expression of *CD38* among 14 subclones. (E) Expression of *CD38* among 14 subclones. Statistical test: Mann Whitney U test in (E): Log2(CPM+1). Significance level: * p<0.05, ** p<0.01, *** p<0.001.

Intra-patient expression heterogeneity was seen, including a t(4;14) patient sample which was obtained after the second relapse and had undergone MM immunotherapy treatment including anti-BCMA (belantamab mafadotin), anti-CD319/SLAMF7 (elotuzumab) and anti-CD38 (daratumumab) therapies (**Supplementary Figure 8**). When checking the expression level of t(4;14) candidates among subclones, we observed cells expressing targets such as *TNFRSF17*/BCMA and *SLAMF7*/CD319 (**Supplementary Figure 8**) indicating gene expression was not responsible for relapse. Other potential new targets were also expressed evenly across all subclones. However, some existing targets did not show good coverage over all subclones. For instance, even though *CD38* was ubiquitously expressed across 49 patient samples (96%), its expression was significantly depleted in several subclones in the anti-CD38 treated sample (p<0.001, **Figure 4D-E**). Taken together, this indicated a mechanism of antigen escape in some subclones which could result in immune-therapy resistance.

We also noticed from the single-cell data that some surface proteins that were not selected as candidates due to insufficient expression in bulk RNA-seq data could be highly expressed in some subclones. For instance, *CHRM3* was highly expressed in only three subclones whereas it was not selected as a candidate due to insufficient expression across all cells (**Supplementary Figure 8**) but has minimal toxicity. This shows that combinations of therapies against multiple targets could be used to fully cover all subclones.

### Alternative splicing as a mechanism of antigen escape

Aberrant RNA splicing was previously found in patient samples following CD19 CAR-T cell therapy, where loss of the epitope through alternative splicing was identified in B-ALL (7). To investigate the possibility of this in MM we performed alternative splicing (AS) analysis on the identified candidate targets both at the population and subgroup levels (**Figure 5A-B and Supplementary Figure 9, Supplementary Table 5**).

**Figure 5.**
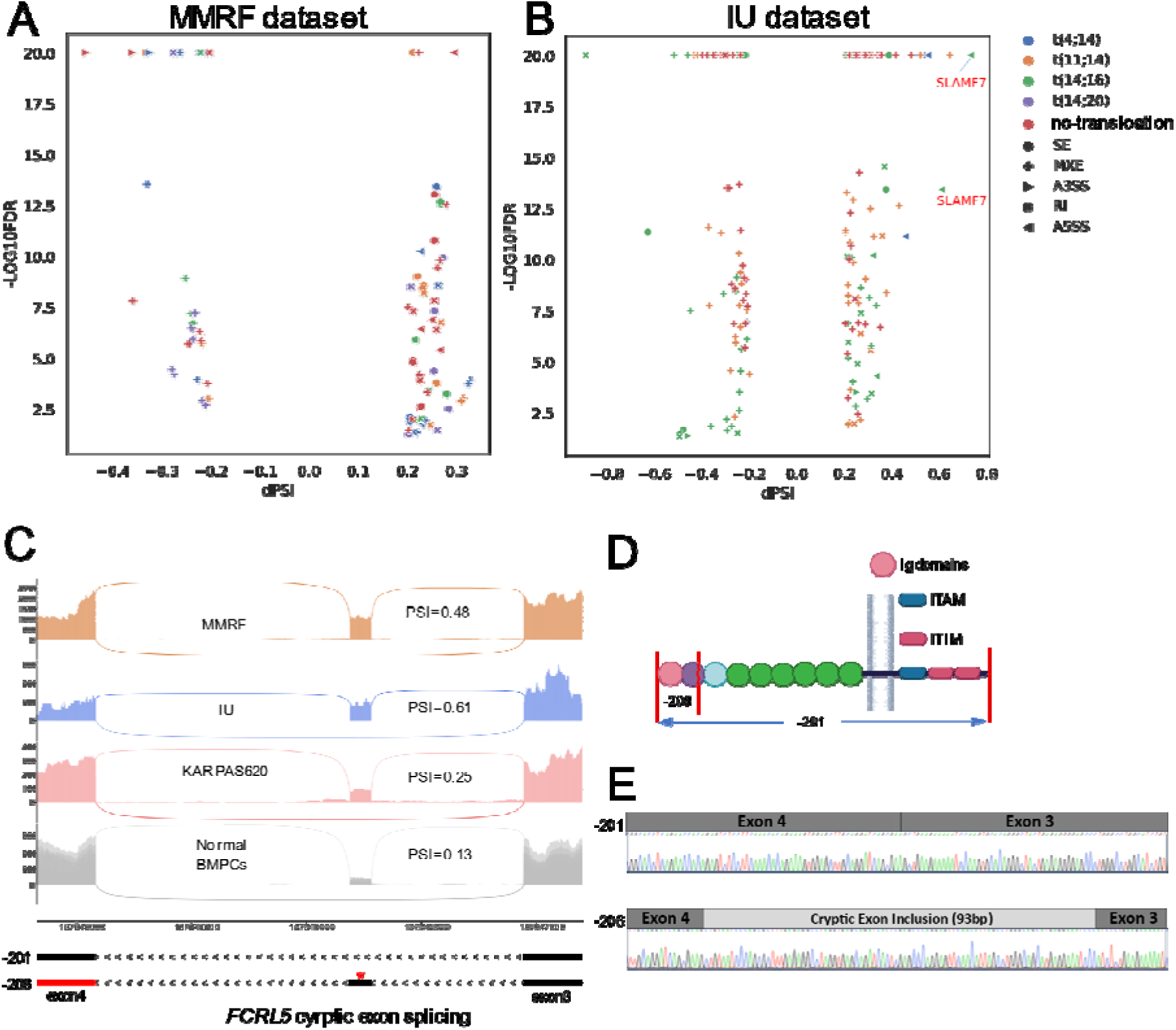
Impact of aberrant splicing towards candidate target gene expression. Summary of most significant alternative splicing events identified from primary (A) and high-risk (B) subtypes, respectively. (C) Sashimi plots indicate inclusion levels of a cryptic exon in *FCRL5* in different samples. Highlighted star: in-frame stop codon. −201, −202, −203, −206: transcript variants of *FCRL5* defined in Ensembl. (D) A schematic plot indicating the truncated domains of a *FCRL5*-206 encoded protein. (E) The cryptic exon validated by RT-PCR and Sanger sequencing.

Interesting alternative splicing events identified in *SLAMF7/CD319* included the previously reported mutually exclusive exon (MXE) event in t(14;16) samples, which results in loss of the transmembrane domain and a secreted product (49).

An alternative splicing event was also identified in *FCRL5*/FCrH5 in both datasets (**Figure 5C-D**), which was further verified by RT-PCR and Sanger sequencing (**Figure 5E**, **Supplementary Figure 9**). In this case, a cryptic exon between exons 3 and 4 was present at higher levels in myeloma samples (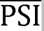=0.24 and 0.21 for MMRF and IU datasets, respectively) than in normal bone marrow plasma cells (BMPC) (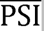=0.13). Inclusion of the cryptic exon is predicted to lead to the switch from a coding transcript to a truncated protein after the first Ig domain (**Figure 5C-D**). Although these patients had not been treated with anti-FCrH5, the presence of an AS event in treatment-naïve patients may lead to the selection of this variant in treated patients as a mechanism of therapy resistance.

## Discussion

The transcriptomic landscape of MM can be used to identify key genes that are expressed on the cell surface and could be targeted by immunotherapies. Several targets for MM already exist including BCMA, CD38, GPRC5D, CD319, and FcRH5 and although treatment of MM patients with immunotherapies targeting these proteins has been successful, patients still relapse through tumor extrinsic (T cell exhaustion) or intrinsic (deletion/mutation) mechanisms due to tumor heterogeneity. Therefore, identifying more potential targets on MM cells could increase the arsenal to attack these cells at subsequent relapses.

Ideally, large proteomic datasets would be available to identify new targets but these are few and far between (50). Several groups have used proteomic techniques to identify and validate new targets (14, 51, 52), but the vast wealth of data exists in RNA expression studies and associated genomic information making them a good starting place to identify novel targets.

Here we have used two genomic and transcriptomic datasets, in combination with external databases, to capture potential immunotherapeutic targets which are expressed on the surface of myeloma cells but not other tissues. Using the genomic information available we determined if there were unique targets specific to key myeloma subgroups, including high-risk subgroups such as del(17p) and gain 1q. Although it would be easier to use uniform targets across all MM patients, there may be some benefit in identifying and targeting subgroup-specific targets, especially those in high-risk groups. Subgroup-specific targets may also be more uniquely expressed on MM cells compared to normal tissues, allowing safer treatment of patients. We also provide a resource for other researchers to further interrogate our data to aid in the further identification of targets in combination with future large proteomic datasets.

Although our analysis is primarily bioinformatic in nature we were able to validate over-expression of genes in subgroups by flow cytometry. Several targets were only expressed in cell lines with specific translocations, for example ROBO3 is expressed in most cell lines except those with a t(11;14) or CD109 in t(4;14) samples. For high-risk groups, we were able to identify genes upregulated in these subgroups and validate them using CRISPR-Cas9 knock out models and flow cytometry. Patients with biallelic *TP53* abnormalities had upregulation of ADAM28, which we confirmed by knockout of *TP53* and flow cytometry. Moreover, we demonstrated the capacity of flow cytometry for precise surface protein detection, which is essential in discerning surface phenotypes of cells. Flow cytometric analysis of cell surface targets may be a better method to determine which antigens could be clinically targeted, due to its current widespread use in diagnostic laboratories.

Antigen escape is a recognized mechanism of therapy resistance in patients treated with anti-BCMA or anti-GPRC5D, and mainly happens through deletion or mutation of the target gene (53, 54). In other B cell malignancies alternative splicing has been described as an additional mechanism of resistance where the epitope recognized by a CAR-T against CD19 was spliced out (55). Here we found a potential intrinsic mechanism of resistance to anti-FcRH5 therapies, where a native splice variant of *FCRL5* results in inclusion of a cryptic exon which would result in loss of the translated protein. This splice variant was validated by RT-PCR in a treatment naïve patient sample, and so could be a mechanism of resistance to be monitored in the future.

In conclusion, we have identified several novel potential immunotherapeutic targets both across all myeloma patients and within genomic subgroups and validated their expression on the surface of MM cells. This is a step towards precision medicine in immunotherapy where patients could have treatments specific to their proteomic profiles which is based on genomic profiling.

## Materials and Methods

### RNA-seq data preprocessing

Bulk RNA-sequencing from 931 CD138+ sorted bone marrow samples were used, including 837 (754 newly diagnosed and 83 relapsed) samples from the MMRF CoMMpass study (MMRF IA18), and 94 (44 newly diagnosed and 50 relapsed) samples generated in-house (IU dataset, **Table 1**). In-house samples were collected from the Indiana Myeloma Registry, a prospective, non-interventional, observational study (NCT03616483) where patients gave informed consent for use of samples for research purposes. The study was approved by Indiana University IRB (#1804208190). Eight samples from normal bone marrow plasma cells (normal dataset) were obtained from previous studies (GSE110486 & GSE114816). Reads were aligned to the hg38 reference genome with gene annotation from Gencode database (V35) using STAR. Transcript-level expression was measured by transcript-per-million (TPM) using Salmon (Quasi-mode mapping, ‘validate map’ mode), aggregated up to gene-level expression, and converted to log_2_(TPM+1) to stabilize variance. Quantile normalization was subsequently conducted for 19,892 protein coding genes to remove batch effects.

Bulk RNA-seq data of 19 MM cell lines from DepMap and an isogenic MM cell line (AMO1) with and without biallelic *TP53* alterations engineered into it (GSE132340) (56) were pre-processed in the same way.

### Differential expression analysis

Differential gene expression analysis was conducted using LIMMA. Significantly differentially expressed genes (DEGs) were further identified as |Log_2_ fold-change|>0.485, log_2_(TPM+1)>1 and FDR<0.05. Pair-wise differential analysis was conducted using either a Mann-Whitney U test (N>3) or T-test (N≤3) with a significance level of p<0.05.

### Identifying high-confidence surface proteins expressed in MM

Expressed surface proteins were obtained from five databases/studies: The Human Protein Atlas (57) (n=1,703); SURFY (58) (n=2,889); Town et al. (59) (n=4,396), Cunha et al. (60) (n=3,744) and Diaz-Ramos et al. (61) (n=1,090). Each gene encoding a cell surface protein was scored between 0 and 5, indicating its cumulative occurrence in these databases/studies. Proteins defined at the plasma membrane, vesicle, cell junctions and focal adhesion sites were annotated as cell surface proteins. Proteins which were present in ≥ three databases were further selected as high-confidence surface proteins. Moreover, we applied the elbow test to the protein expression profile of 2,956 high-confidence surface proteins identified from *in vitro* surface biotinylation proteomic experiments of MM cells (8), which led to the addition of a further 122 proteins. An expression cut-off was determined by examining the RNA expression of 19,783 normalized protein coding genes across the two datasets and selecting the highest 75 percentile, which equated to a log_2_(TPM+1)>3.

### Identifying surface proteins with low side effects

Gene expression levels of surface proteins in 37 healthy tissue types were obtained from GTEx(62) annotation from The Human Protein Atlas. Normalized transcript expression levels for each gene (nTPM) was calculated and a nTPM >50 was used as the cut-off for differentiating high RNA expression of surface proteins in any healthy tissue. Genes encoding surface proteins that were highly expressed in MM but present in fewer than three healthy tissues were defined as candidates with low tissue side effects. Immune-related organs were excluded from the analysis and instead gene expression levels in cells from whole blood were calculated from The Human Protein Atlas v21 which includes 19 blood cell types, including lymphoid and myeloid compartments. The maximum gene expression in myeloid compartment cells (basophils, classical monocytes, eosinophil, intermediate monocytes, myeloid dendritic cells, neutrophils, non-classical monocytes, and plasmacytoid dendritic cells) was used to indicate potential toxicity to blood cells.

### Prioritizing candidates

Candidates identified at the population level were prioritized by their median expression level in the NDMM MMRF dataset. Candidates identified at the subtype-level were organized by their occurrence (from more to less), clustered by their expression similarities (Euclidean distances), and prioritized for the specific subgroup if their median expression level was significantly higher than the population median (Log_2_FC>0.5).

### Pathway analysis

Over-representation analysis was conducted for any given gene list against Kyoto Encyclopedia of Genes and Genome (KEGG) pathways and Gene Ontology (GO) biological processes.

### Survival analysis

Overall survival (OS) and progression-free survival (PFS) was conducted for NDMM patients in the MMRF dataset. The Cox proportional hazard model was used to calculate the hazard ratio for given variables. The Log-rank test was used to measure the survival difference between groups measured in Kaplan-Meier plots. P=0.05 was used as the significance level cut-off in both statistical tests.

Within each molecular subgroup, patients were stratified into four equal groups (quartiles) based on the expression levels of given genes. Survival analyses were performed to compare patient outcomes across these quartiles. t(14;16) and t(14;20) subgroups were combined into a t(MAF) subgroup for the survival analysis to increase the statistical power.

### Differential RNA splicing analysis

rMATS was used to identify differential splicing events between designated MM subgroups and the normal samples. Percent spliced-in (PSI) was used to measure the splicing level of alternative splicing (AS) events while deltaPSI (dPSI) was used to measure the average splicing difference between two groups. AS events with |dPSI|>20%, FDR<0.05 (likelihood ratio test) and more than 10 junction reads detected in >25% samples were subsequently identified as significant AS events.

### Preprocessing and peak-calling of ChIP-seq data

ChIP-seq data from T cells (anti-MAF) (GSE72266) were aligned to hg38 using BWA, PCR duplicates marked by Picard, peaks called using MACS2, and ranked by their FDR. Previously defined chromatin states in seven MM patient-derived xenograft samples were used to define MM super-enhancers (63).

### Gene essentiality

Gene essentiality scores (CERES) of 18 MM cell lines were obtained for DepMap. A CERES score=0 indicates no essentiality (no effect on cell growth with gene knock-out) while a CERES score of 1 indicates a strong essentiality (lack of growth after knock-out). Genomic abnormalities of cell lines were obtained from a previous annotation (64). 1q status was defined by *CKS1B* copy number from the DepMap annotation. Biallelic/monoallelic *TP53* annotations were previously described (49).

### Single-cell data processing and analysis

49 single-cell RNA-seq samples from a previous study (47) were utilized, including 35 patient samples that overlap with this study. We followed the previously defined methods (47) for data preprocessing, gene/sample filtering and expression-level normalization. Counts per million (CPM) and Log_2_(CPM+1) were used to measure gene expression. Subclones were identified using the build-in clustering functions for in Seurat(65). A gene was defined as expressed in a subclone when >25% of the cells expressed the gene.

### Cell lines culture

Myeloma cell lines were grown in suspension in RPMI-1640 (Gibco) supplemented with 10% fetal bovine serum and 1% Penicillin-streptomycin (Gibco). All cells were incubated at 37°C, 5% CO_2_. Cell lines were tested for mycoplasma using MycoAlert Mycoplasma Detection Kit (Lonza) and were negative.

### TP53 knockout using CRISPR-Cas9 technology

AMO1 *TP53* knockout cells were generated using CRISPR-Cas9 technology. Ribonucleoproteins (RNPs) consisting of Cas-9 and gRNAs (**Supplementary Table 1**) were assembled to target *TP53* in the AMO1 cell line which has two functional copies. After single-cell cloning by limiting dilution, cells were tested for *TP53* status by PCR and Sanger sequencing as well as Western blot.

### Flow cytometry

Flow-cytometric analysis was performed to examine the extracellular expression of BCMA, CD49d, ADAM28, ROBO3 or CD109 in different myeloma cell lines. 1×10^6^ cells were harvested, washed with PBS, and resuspended in 100 µL of cell staining buffer (Biolegend) and Fc blocked using Human TruStain FcX (Biolegend) for 10 mins at room temperature to reduce non-specific immunofluorescent staining. Cells were either directly or indirectly stained with conjugated antibodies. For the direct immunostaining, cells were labeled with PE anti-human BCMA (Biolegend), PE anti-human CD49d (BD Biosciences) or Alexa Fluor 647 anti-human ADAM28 (Biotechne) conjugated antibody or with the matching mouse isotype immunoglobulin controls, PE mouse IgG2a (Biolegend), PE mouse IgG1 (Invitrogen) or Alexa Fluor 647 mouse IgG1 (Biotechne), respectively for 30 mins on ice. For indirect immunostaining, cells were incubated with the unconjugated anti-human CD109 (Invitrogen) or anti-human ROBO3 (Invitrogen) or with the matching isotype, mouse IgG1 (Invitrogen) or goat IgG (Biotechne), respectively, for 1 hour at room temperature. Cells were incubated with PE-conjugated goat anti-mouse IgG (Invitrogen) or PE-conjugated donkey anti-goat (Biotechne) secondary antibodies at room temperature for 30 minutes. After staining, cells were washed, resuspended in the staining buffer and analyzed by flow cytometry. 10000 events per sample were acquired using BD LSRFortessa X-20 Cell Analyzer (BD Bioscience, San Jose, CA). Data were analyzed by FlowJo v10.4 (TreeStar, Palo Alto, CA).

### Immunoblotting analysis

Cells were lysed using RIPA buffer (150 mM NaCl, 50 mM TrisHCl (pH 7.4), 1% NP40, 0.5% sodium deoxycholate, 0.1% SDS, 1 mM EDTA, 10 mM, NaF; pH 8.0) and a cocktail of protease inhibitors (cOmplete, Roche) and phosphatase inhibitors (Pierce tablets, Thermofisher Scientific). Cell lysates were incubated on ice for 20 mins and centrifuged at 16,000 xg for 20 mins. Supernatants were collected and protein concentrations were measured with protein Broad range quantification Kit (Thermofisher) using Qubit 4 Fluorometer. 40 µg of protein was loaded on Mini-PROTEAN TGX Precast (Biorad) SDS-PAGE gels followed by transfer to nitrocellulose membranes. After blocking with non-fat dry milk for 1h at room temperature, membranes were probed at 4 °C with the following primary antibodies overnight: anti-p53 (Santa Cruz Biotechnology), or anti-β-actin (Cell signaling). Membranes were incubated with the HRP-conjugated secondary antibodies anti-rabbit (ThermoFisher Scientific). Immune complexes were detected by chemical enhanced luminescence ECL using SuperSignal West Pico Plus Chemiluminescent Substrate (ThermoFisher Scientific). Images were acquired using ChemiDoc Imaging system (Biorad). Densitometric analysis to determine protein expression was performed using Image Lab software (Version 6.0.1, Biorad).

## Author Contributions

E.L. analyzed data and wrote the paper. O.J. and R.S. developed knockout cell lines, performed western blots and flow cytometry. T.S.J. analyzed single cell data. V.C., F.Z. and H.H. provided resources, C.P. provided protocols and critical feedback, R.A. and A.S. provided clinical samples and input. B.A.W. analyzed data, provided resources, and wrote the paper.

## Acknowledgements

The Indiana Myeloma Registry is funded in part by support from the Indiana University Precision Health Initiative, Miles for Myeloma, the Harry and Edith Gladstein Chair, and the Omar Barham Fighting Cancer Fund. Computational infrastructure at Indiana University (Indianapolis, IN) was funded in part by Lilly Endowment Inc. through the Indiana University Pervasive Technology Institute. We thank the Center for Medical Genomics at the Indiana University School of Medicine, a core supported by the NCI Cancer Center P30 support grant CA082709, for their expertise in carrying out the sequencing of patient samples. This study used the Multiple Myeloma Research Foundation (MMRF) CoMMpass Dataset. The authors acknowledge the efforts of the MMRF research consortium to provide the fundamental resource for our study.

R.A., B.A.W., and M.A.Z. received research support from Genentech to carry out this study. B.A.W. is partly funded by the Daniel and Lori Efroymson Chair. E.L. received a research fellowship grant from the Multiple Myeloma Research Foundation. T.S.J. received a research fellowship grant from the Multiple Myeloma Research Foundation and is partly funded by the Agnes Beaudry Investigator in Myeloma Research fund.

## Data availability

Genomic data generated in this study have been submitted to dbGaP with accession number phs003772.v1.p1. Previous datasets including the MMRF RNA-seq data (accession number phs000748) and Indiana University single-cell data (accession number phs003220.v2) are available at dbGaP. ChIP-seq data from T cells (anti-MAF) can be obtained from gene expression omnibus (GEO) database (accession ID GSE72266). RNA-seq data of *TP53* knockout/WT in AMO1 cell lines can be obtained from GEO accession GSE132340. RNA-seq data of normal bone marrow plasma cell samples can be obtained from GEO accession GSE110486 and GSE114816.

