## Supplementary Data for "Utilizing Genomics to Identify Novel Immunotherapeutic Targets in Multiple Myeloma High-Risk Subgroups"

**Affiliations:**

### Supplementary Data

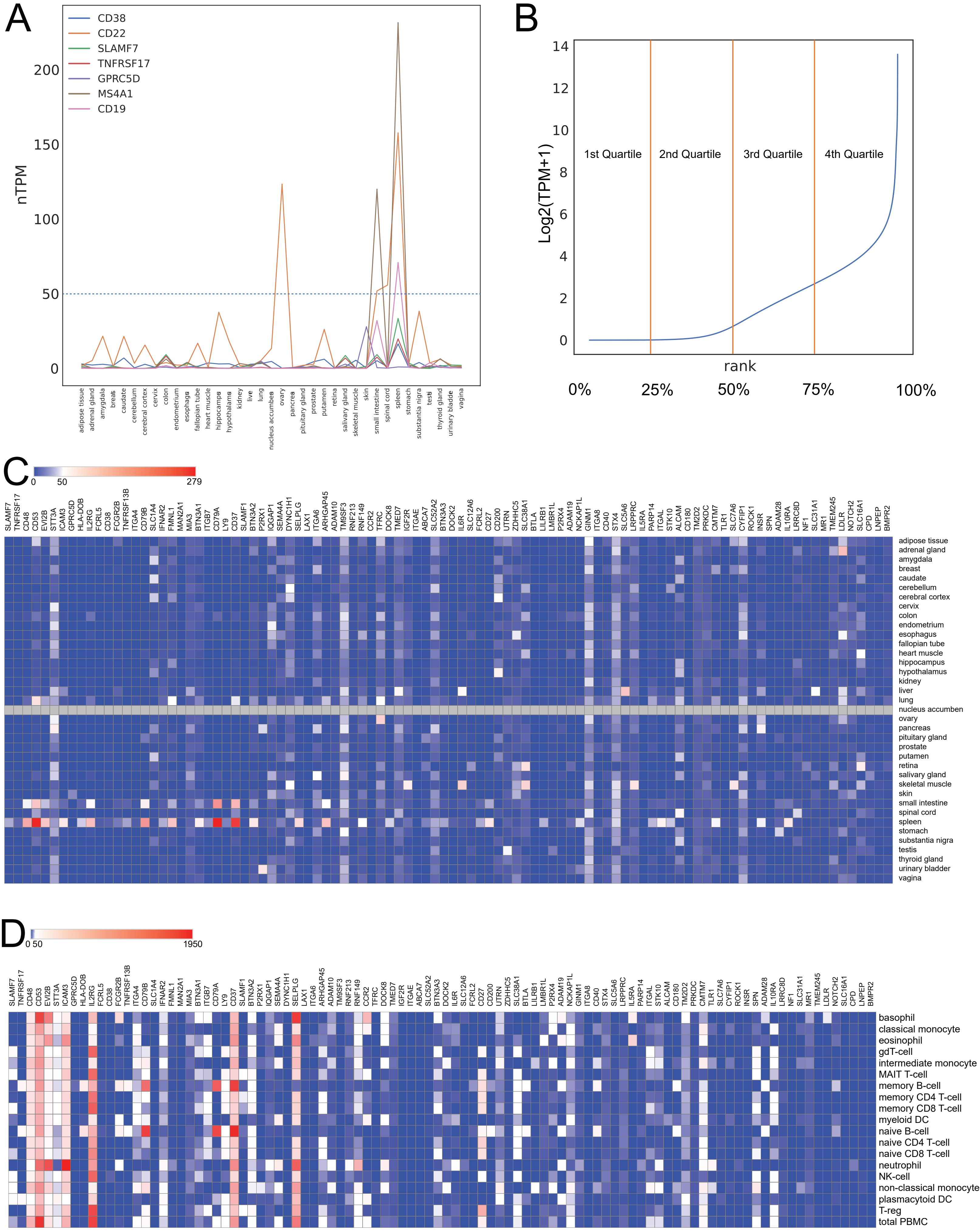

A
CD38
CD22
SLAMF7
TNFRSF17
GPRC5D
MS4A1
CD19
nTPM
0
50
100
150
200
adipose tissue
adrenal gland
amygdala
breas
caudate
cerebellum
cerebral cortex
cervix
colon
endometrium
esophagus
fallopian tube
heart muscle
hippocampus
hypothalamus
kidney
live
lung
nucleus accumbens
ovary
pancreas
pituitary gland
prostate
putamen
retina
salivary gland
skeletal muscle
skin
small intestine
spinal cord
spleen
stomach
substantia nigra
testi
thyroid gland
urinary bladde
vagina
B
Log2(TPM+1)
0
2
4
6
8
10
12
14
1st Quartile
2nd Quartile
3rd Quartile
4th Quartile
rank
0%
25%
50%
75%
100%
C
0
50
279
SLAMF7 TNFRSF17 CD48 CD53 EVI2B STT3A ICAM3 GPRC5D HLA-DOB IL2RG FCRL5 CD38 FCGR2B TNFRSF13B ITGA4 CD79B SLC1A4 IFNAR2 FMNL1 MAN2A1 MIA3 BTN3A1 ITGB7 CD79A LY9 CD37 SLAMF1 BTN3A2 P2RX1 IQGAP1 SEMA4A DYNC1H1 SELPLG LAX1 ITGA6 ARHGAP45 ADAM10 TM9SF3 RNF213 RNF149 CCR2 TFRC DOCK8 TMED7 IGF2R ITGAE ABCA7 SLC52A2 BTN3A3 DOCK2 IL6R SLC12A6 FCRL2 CD27 CD200 UTRN ZDHHC5 SLC38A1 BTLA LILRB1 LMBR1L P2RX4 ADAM19 NCKAP1L GINM1 ITGA8 CD40 STX4 SLC5A6 LRPPRC IL5RA PARP14 ITGAL STK10 ALCAM CD180 TM2D2 PRKDC CMTM7 TLR1 SLC7A6 CYFIP1 ROCK1 INSR SPN ADAM28 IL10RA LRRC8D NF1 SLC31A1 MR1 TMEM245 LDLR NOTCH2 SLC16A1 CPD LNPEP BMPR2
adipose tissue
adrenal gland
amygdala
breast
caudate
cerebellum
cerebral cortex
cervix
colon
endometrium
esophagus
fallopian tube
heart muscle
hippocampus
hypothalamus
kidney
liver
lung
nucleus accumben
ovary
pancreas
pituitary gland
prostate
putamen
retina
salivary gland
skeletal muscle
skin
small intestine
spinal cord
spleen
stomach
substantia nigra
testis
thyroid gland
urinary bladder
vagina
D
0 50
1950
SLAMF7 TNFRSF17 CD48 CD53 EVI2B STT3A ICAM3 GPRC5D HLA-DOB IL2RG FCRL5 CD38 FCGR2B TNFRSF13B ITGA4 CD79B SLC1A4 IFNAR2 FMNL1 MAN2A1 MIA3 BTN3A1 ITGB7 CD79A LY9 CD37 SLAMF1 BTN3A2 P2RX1 IQGAP1 SEMA4A DYNC1H1 SELPLG LAX1 ITGA6 ARHGAP45 ADAM10 TM9SF3 RNF213 RNF149 CCR2 TFRC DOCK8 TMED7 IGF2R ITGAE ABCA7 SLC52A2 BTN3A3 DOCK2 IL6R SLC12A6 FCRL2 CD27 CD200 UTRN ZDHHC5 SLC38A1 BTLA LILRB1 LMBR1L P2RX4 ADAM19 NCKAP1L GINM1 ITGA8 CD40 STX4 SLC5A6 LRPPRC IL5RA PARP14 ITGAL STK10 ALCAM CD180 TM2D2 PRKDC CMTM7 TLR1 SLC7A6 CYFIP1 ROCK1 INSR SPN ADAM28 IL10RA LRRC8D NF1 SLC31A1 MR1 TMEM245 LDLR NOTCH2 SLC16A1 CPD LNPEP BMPR2
basophil
classical monocyte
eosinophil
gdT-cell
intermediate monocyte
MAIT T-cell
memory B-cell
memory CD4 T-cell
memory CD8 T-cell
myeloid DC
naive B-cell
naive CD4 T-cell
naive CD8 T-cell
neutrophil
NK-cell
non-classical monocyte
plasmacytoid DC
T-reg
total PBMC

**Supplementary Figure 1. Characteristics of candidate targets identified from MM populations.**

**A**. RNA expression level of current CAR-T cell targets of blood cancers in healthy organs documented in the human protein atlas (THPA). **B**. Distribution of mRNA expression of 19,892 protein coding genes in MMRF and IU cohort. **C**. Expression level of 98 candidates in healthy organs documented in the human protein atlas (THPA). **D**. Expression level of 98 candidates in myeloid blood cells documented in 'The Human Protein Atlas' database.

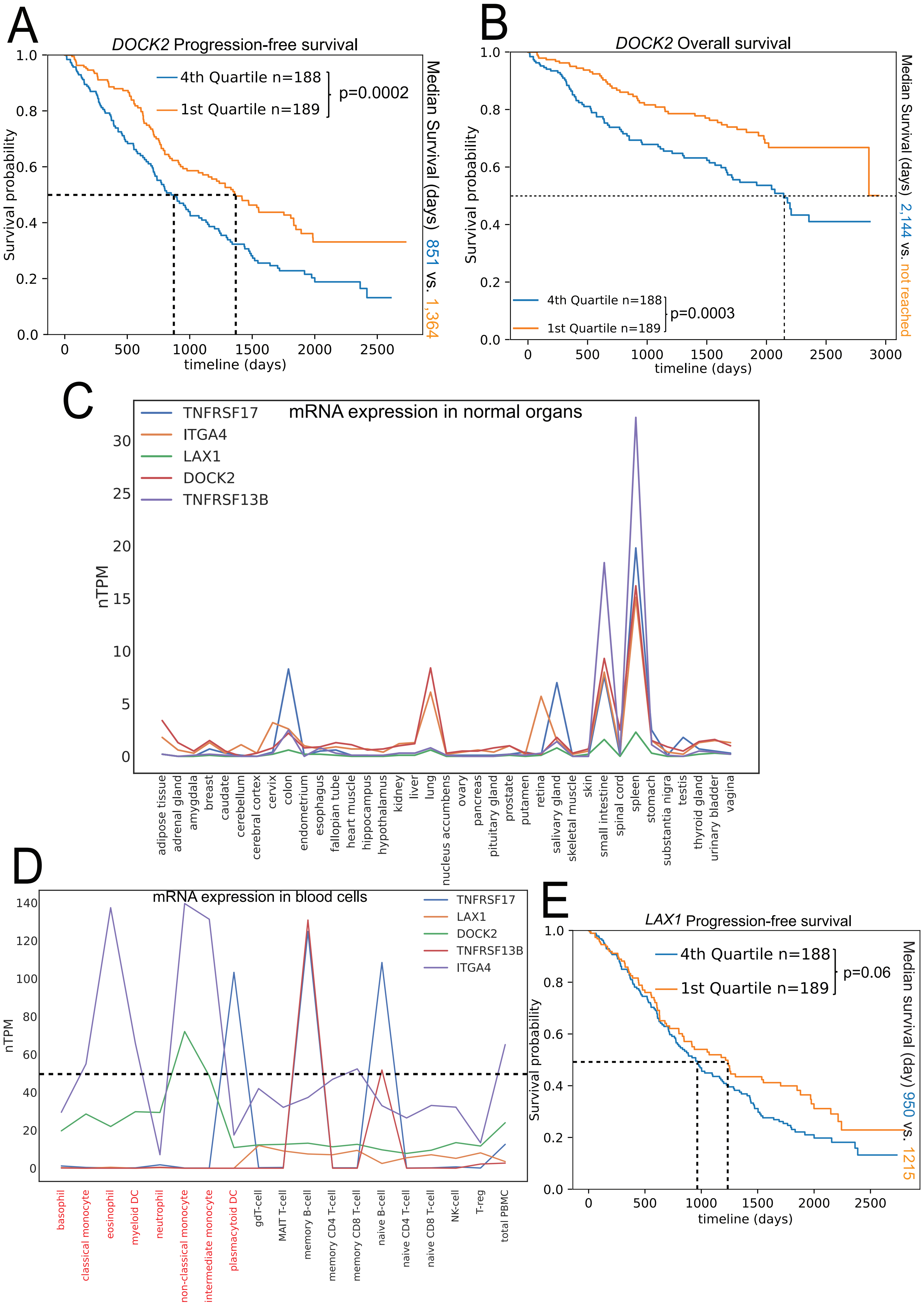

A
DOCK2 Progression-free survival
4th Quartile n=188
1st Quartile n=189
p=0.0002
Median Survival (days) 851 vs. 1,364
Survival probability
timeline (days)
B
DOCK2 Overall survival
4th Quartile n=188
1st Quartile n=189
p=0.0003
Median Survival (days) 2,144 vs. not reached
Survival probability
timeline (days)
C
mRNA expression in normal organs
TNFRSF17
ITGA4
LAX1
DOCK2
TNFRSF13B
nTPM
adipose tissue
adrenal gland
amygdala
breast
caudate
cerebellum
cerebral cortex
cervix
colon
endometrium
esophagus
fallopian tube
heart muscle
hippocampus
hypothalamus
kidney
liver
lung
nucleus accumbens
ovary
pancreas
pituitary gland
prostate
putamen
retina
salivary gland
skeletal muscle
skin
small intestine
spinal cord
spleen
stomach
substantia nigra
testis
thyroid gland
urinary bladder
vagina
D
mRNA expression in blood cells
TNFRSF17
LAX1
DOCK2
TNFRSF13B
ITGA4
nTPM
basophil
classical monocyte
eosinophil
myeloid DC
neutrophil
non-classical monocyte
intermediate monocyte
plasmacytoid DC
gdT-cell
MAIT T-cell
memory B-cell
memory CD4 T-cell
memory CD8 T-cell
naive B-cell
naive CD4 T-cell
naive CD8 T-cell
NK-cell
T-reg
total PBMC
E
LAX1 Progression-free survival
4th Quartile n=188
1st Quartile n=189
p=0.06
Median survival (day) 950 vs. 1215
Survival probability
timeline (days)

**Supplementary Figure 2. Toxicity and association with patient survival of identified population-based candidates.**

**A**. Progression-free survival of patients with different *DOCK2* expression levels. **B**. Overall survival of patients with different *DOCK2* expression levels. **C**. Expression level of existing targets and selected candidates in healthy organs. **D**. Expression level of existing targets and selected candidates in blood cells. **E**. Progression-free survival of patients with different *LAX1* expression. Statistical test in Kaplan-Meier curves: Log-rank test. Highlighted labels in **D**: myeloid blood cells.

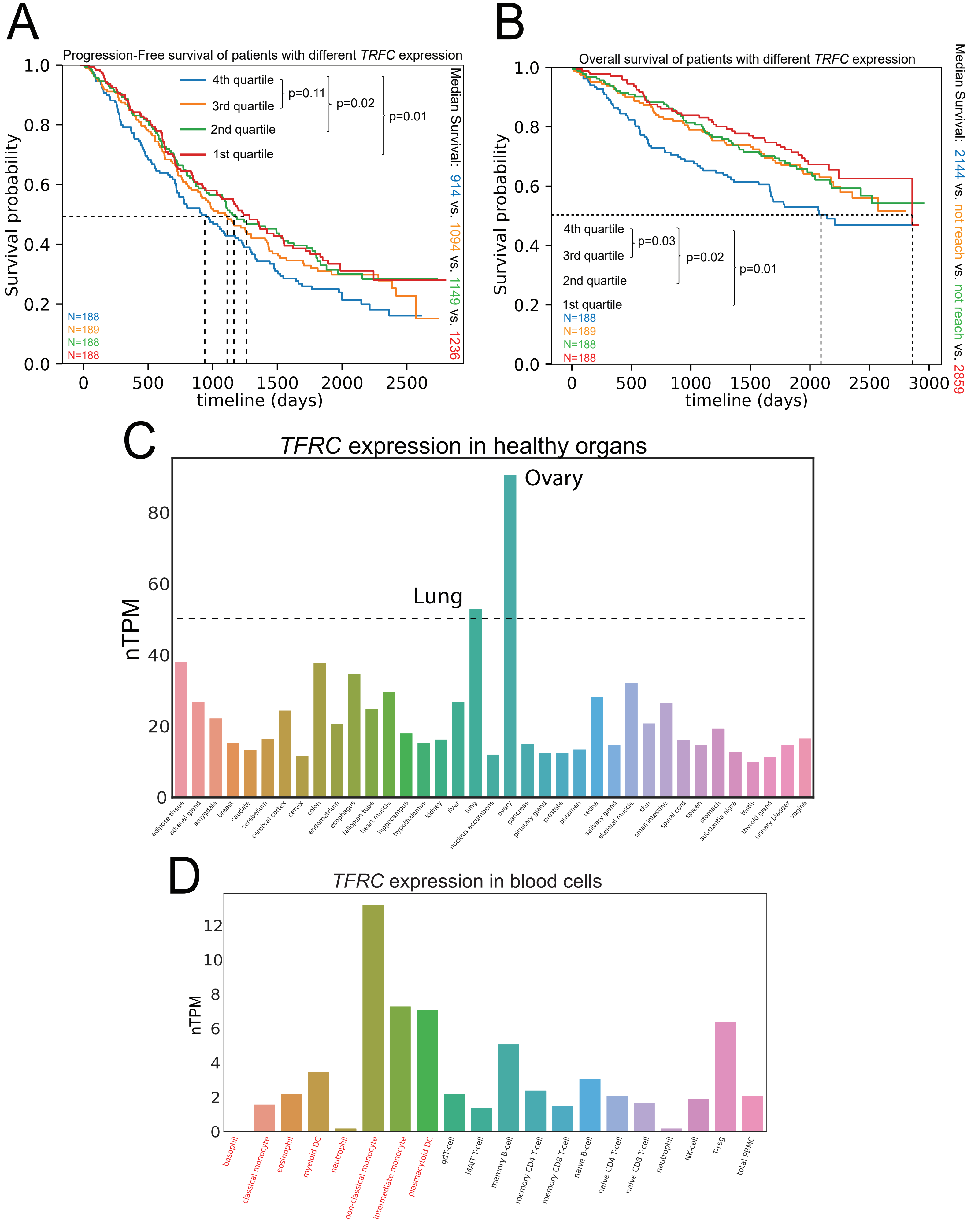
A
Progression-Free survival of patients with different TRFC expression
4th quartile
3rd quartile
2nd quartile
1st quartile
p=0.11
p=0.02
p=0.01
Median Survival: 914 vs. 1094 vs. 1149 vs. 1236
N=188
N=189
N=188
N=188
Survival probability
timeline (days)
B
Overall survival of patients with different TRFC expression
4th quartile
3rd quartile
2nd quartile
1st quartile
p=0.03
p=0.02
p=0.01
N=188
N=189
N=188
N=188
Median Survival: 2144 vs. not reach vs. not reach vs. 2859
Survival probability
timeline (days)
C
TFRC expression in healthy organs
Ovary
Lung
nTPM
adipose tissue
adrenal gland
amygdala
breast
caudate
cerebellum
cerebral cortex
cervix
colon
endometrium
esophagus
fallopian tube
heart muscle
hippocampus
hypothalamus
kidney
liver
lung
nucleus accumbens
ovary
pancreas
pituitary gland
prostate
putamen
retina
salivary gland
skeletal muscle
skin
small intestine
spinal cord
spleen
stomach
substantia nigra
testis
thyroid gland
urinary bladder
vagina
D
TFRC expression in blood cells
nTPM
basophil
classical monocyte
eosinophil
myeloid DC
neutrophil
non-classical monocyte
intermediate monocyte
plasmacytoid DC
gdT-cell
MAIT T-cell
memory B-cell
memory CD4 T-cell
memory CD8 T-cell
naive B-cell
naive CD4 T-cell
naive CD8 T-cell
neutrophil
NK-cell
T-reg
total PBMC

**Supplementary Figure 3. Characteristics of a highly essential candidate *TRFC***

**A**. Progression-free survival of patients with different *TFRC* expression levels. **B**. Overall survival of patients with different *TFRC* expression levels. **C**. *TFRC* expression level in healthy organs. **D**. *TFRC* expression level in blood cells. Statistical test in Kaplan-Meier curves: Logrank test. Highlighted labels in **D**: myeloid blood cells.

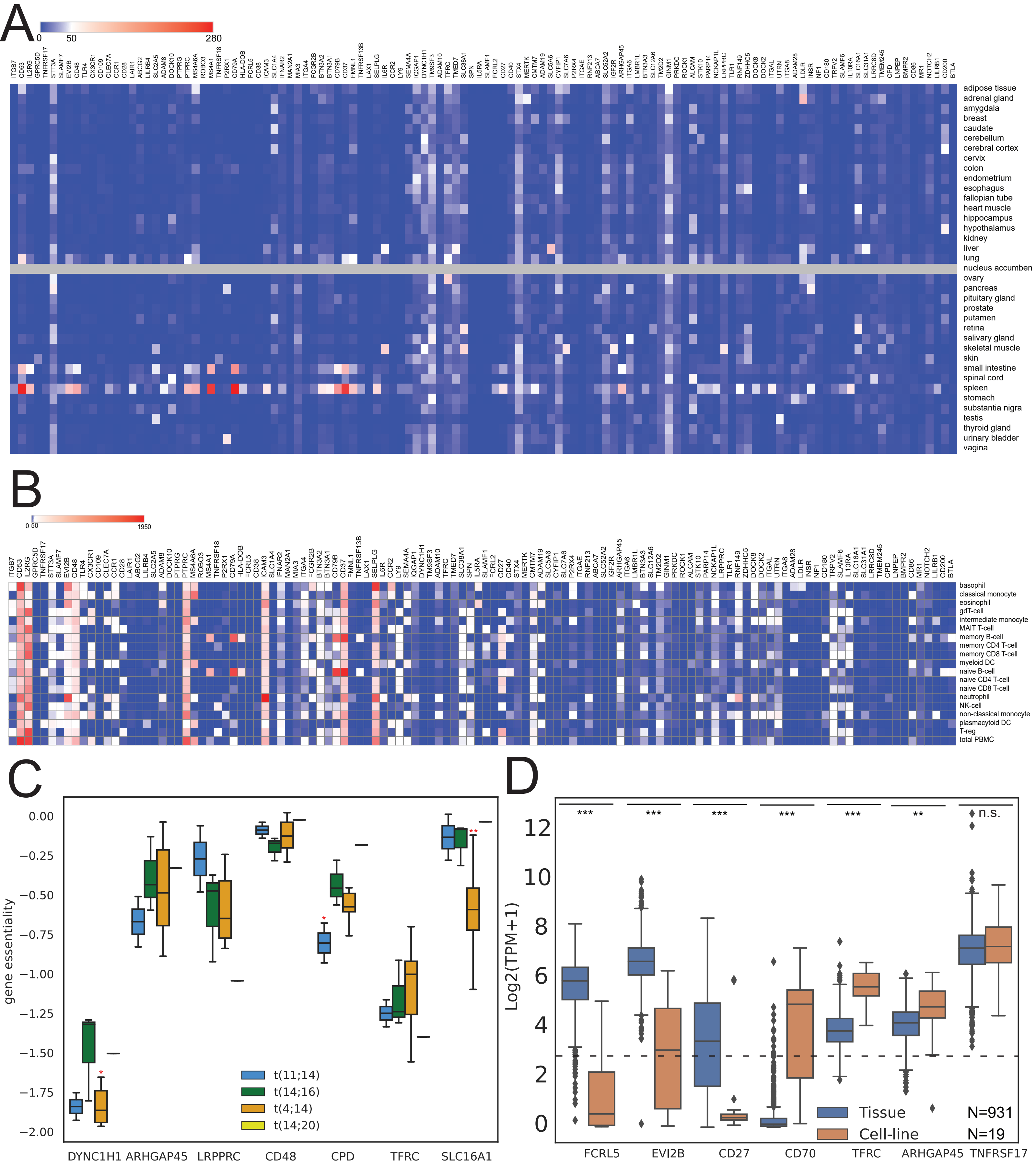
A
0
50
280
ITGB7 CD53 IL2RG GPRC5D TNFRSF17 STT3A SLAMF7 EVI2B CD48 TLR4 CX3CR1 CD109 CLEC7A CCR1 CD28 LAIR1 ABCG2 LILRB4 SLC2A5 ADAM8 DOCK10 PTPRG PTPRC MS4A6A ROBO3 MS4A1 TNFRSF18 P2RX1 CD79A HLA-DOB FCRL5 CD38 ICAM3 SLC1A4 IFNAR2 MAN2A1 MIA3 ITGA4 FCGR2B BTN3A2 BTN3A1 CD79B CD37 FMNL1 TNFRSF13B LAX1 SELPLG IL6R CCR2 LY9 SEMA4A IQGAP1 DYNC1H1 TM9SF3 ADAM10 TFRC TMED7 SLC38A1 SPN IL5RA SLAMF1 FCRL2 CD27 CD40 STX4 MERTK CMTM7 ADAM19 SLC5A6 CYFIP1 SLC7A6 P2RX4 ITGAE RNF213 ABCA7 SLC52A2 IGF2R ARHGAP45 ITGA6 LMBR1L BTN3A3 SLC12A6 TM2D2 GINM1 PRKDC ROCK1 ALCAM STK10 PARP14 NCKAP1L LRPPRC TLR1 RNF149 ZDHHC5 DOCK8 DOCK2 ITGAL UTRN ITGA8 ADAM28 LDLR INSR NF1 CD180 TRPV2 SLAMF6 IL10RA SLC31A1 SLC16A1 LRRC8D TMEM245 CPD LNPEP BMPR2 CD86 MR1 NOTCH2 LILRB1 CD200 BTLA
adipose tissue
adrenal gland
amygdala
breast
caudate
cerebellum
cerebral cortex
cervix
colon
endometrium
esophagus
fallopian tube
heart muscle
hippocampus
hypothalamus
kidney
liver
lung
nucleus accumben
ovary
pancreas
pituitary gland
prostate
putamen
retina
salivary gland
skeletal muscle
skin
small intestine
spinal cord
spleen
stomach
substantia nigra
testis
thyroid gland
urinary bladder
vagina
B
0 50
1950
ITGB7 CD53 IL2RG GPRC5D TNFRSF17 STT3A SLAMF7 EVI2B CD48 TLR4 CX3CR1 CD109 CLEC7A CCR1 CD28 LAIR1 ABCG2 LILRB4 SLC2A5 ADAM8 DOCK10 PTPRG PTPRC MS4A6A ROBO3 MS4A1 TNFRSF18 P2RX1 CD79A HLA-DOB FCRL5 CD38 ICAM3 SLC1A4 IFNAR2 MAN2A1 MIA3 ITGA4 FCGR2B BTN3A2 BTN3A1 CD79B CD37 FMNL1 TNFRSF13B LAX1 SELPLG IL6R CCR2 LY9 SEMA4A IQGAP1 DYNC1H1 TM9SF3 ADAM10 TFRC TMED7 SLC38A1 SPN IL5RA SLAMF1 FCRL2 CD27 CD40 STX4 MERTK CMTM7 ADAM19 SLC5A6 CYFIP1 SLC7A6 P2RX4 ITGAE RNF213 ABCA7 SLC52A2 IGF2R ARHGAP45 ITGA6 LMBR1L BTN3A3 SLC12A6 TM2D2 GINM1 PRKDC ROCK1 ALCAM STK10 PARP14 NCKAP1L LRPPRC TLR1 RNF149 ZDHHC5 DOCK8 DOCK2 ITGAL UTRN ITGA8 ADAM28 LDLR INSR NF1 CD180 TRPV2 SLAMF6 IL10RA SLC31A1 SLC16A1 LRRC8D TMEM245 CPD LNPEP BMPR2 CD86 MR1 NOTCH2 LILRB1 CD200 BTLA
basophil
classical monocyte
eosinophil
gdT-cell
intermediate monocyte
MAIT T-cell
memory B-cell
memory CD4 T-cell
memory CD8 T-cell
myeloid DC
naive B-cell
naive CD4 T-cell
naive CD8 T-cell
neutrophil
NK-cell
non-classical monocyte
plasmacytoid DC
T-reg
total PBMC
C
gene essentiality
0.00
−0.25
−0.50
−0.75
−1.00
−1.25
−1.50
−1.75
−2.00
t(11;14)
t(14;16)
t(4;14)
t(14;20)
DYNC1H1 ARHGAP45 LRPPRC CD48 CPD TFRC SLC16A1
D
Log2(TPM+1)
12
10
8
6
4
2
0
***
***
***
***
***
**
n.s.
Tissue N=931
Cell-line N=19
FCRL5 EVI2B CD27 CD70 TFRC ARHGAP45 TNFRSF17

**Supplementary Figure 4. Characteristics of candidates identified from primary subtypes.** **A**. Expression level of 120 candidates in healthy organs documented in the human protein atlas (THPA). **B**. Expression level of 120 candidates in blood cells documented in 'The Human Protein Atlas' database. **C**. Gene essentiality differences of selected candidates in 18 MM cell-lines from DepMap. **D**. Expression heterogeneity of candidates between tissue and cell-lines. Statistical tests in **C** and **D**: Mann-Whitney U test. Highlighted stars in **C**: significance level of candidates with higher essentiality in their subgroups than average in MM cell-lines. Highlighted stars in **D**: significance level of candidates with differential expression. Significance level: * $p<0.05$; ** $p<0.01$, *** $p<0.001$.

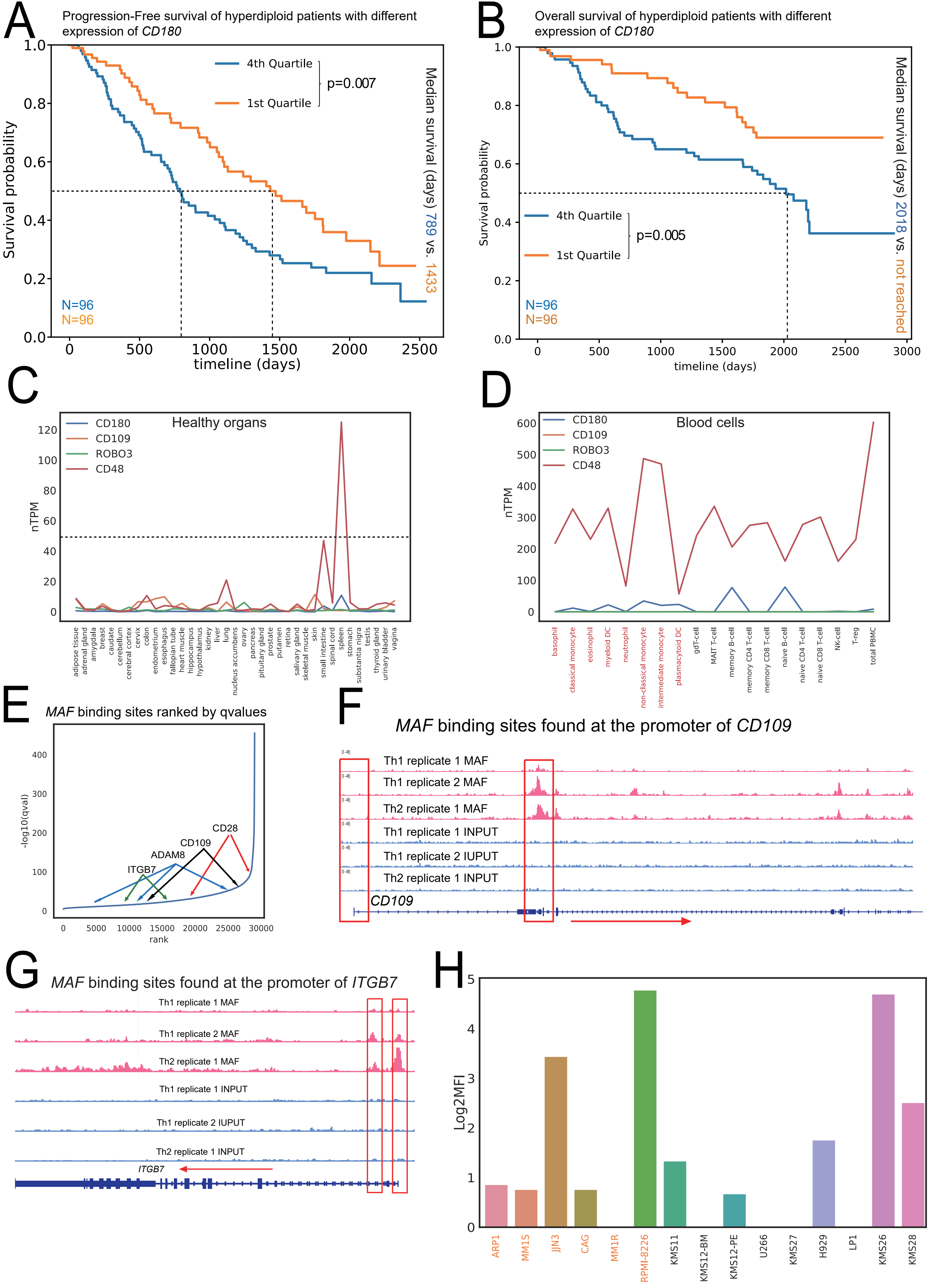
A
Progression-Free survival of hyperdiploid patients with different expression of CD180
4th Quartile
1st Quartile
p=0.007
Survival probability
timeline (days)
Median survival (days) 789 vs. 1433
N=96
N=96
B
Overall survival of hyperdiploid patients with different expression of CD180
4th Quartile
1st Quartile
p=0.005
Survival probability
timeline (days)
Median survival (days) 2018 vs. not reached
N=96
N=96
C
Healthy organs
CD180
CD109
ROBO3
CD48
nTPM
adipose tissue
adrenal gland
amygdala
breast
caudate
cerebellum
cerebral cortex
cervix
colon
endometrium
esophagus
fallopian tube
heart muscle
hippocampus
hypothalamus
kidney
liver
lung
nucleus accumbens
ovary
pancreas
pituitary gland
prostate
putamen
retina
salivary gland
skeletal muscle
skin
small intestine
spinal cord
spleen
stomach
substantia nigra
testis
thyroid gland
urinary bladder
vagina
D
Blood cells
CD180
CD109
ROBO3
CD48
nTPM
basophil
classical monocyte
eosinophil
myeloid DC
neutrophil
non-classical monocyte
intermediate monocyte
plasmacytoid DC
gdT-cell
MAIT T-cell
memory B-cell
memory CD4 T-cell
memory CD8 T-cell
naive B-cell
naive CD4 T-cell
naive CD8 T-cell
NK-cell
T-reg
total PBMC
E
MAF binding sites ranked by qvalues
-log10(qval)
rank
CD28
CD109
ADAM8
ITGB7
F
MAF binding sites found at the promoter of CD109
Th1 replicate 1 MAF
Th1 replicate 2 MAF
Th2 replicate 1 MAF
Th1 replicate 1 INPUT
Th1 replicate 2 IUPUT
Th2 replicate 1 INPUT
CD109
G
MAF binding sites found at the promoter of ITGB7
Th1 replicate 1 MAF
Th1 replicate 2 MAF
Th2 replicate 1 MAF
Th1 replicate 1 INPUT
Th1 replicate 2 IUPUT
Th2 replicate 1 INPUT
ITGB7
H
Log2MFI
ARP1
MM1S
JJN3
CAG
MM1R
RPMI-8226
KMS11
KMS12-BM
KMS12-PE
U266
KMS27
H929
LP1
KMS26
KMS28

**Supplementary Figure 5. Toxicity and association with patient survival of identified subtype-specific candidates.**

**A**. Progression-free survival of hyper-diploid patients with different *CD180* expression levels. **B**. Overall survival of hyper-diploid patients with different *CD180* expression levels. **C**. Selected candidates identified in subtypes and their expression level in healthy organs. **D**. Selected candidates identified in subtypes and their expression level in blood cells. **E**. Four candidates found with *MAF* binding sites from ChIP-seq result. **F**. *MAF* binding sites found at the promoter region of CD109. **G**. *MAF* binding sites found at the promoter region of *ITGB7*. **H**. Protein expression level (Log2MFI) of *CD109* detected by flow cytometry among 15 cell-lines. Statistical test in Kaplan-Meier curves: Logrank test. Highlighted labels in **D**: myeloid blood cells. Highlighted boxes in **F** and **G**: promoter regions. Red arrows: transcription direction. Highlighted cell-lines in **H**: t(14;16) cell-lines. KMS11 is a t(14;16) and t(4;14) cell-line.

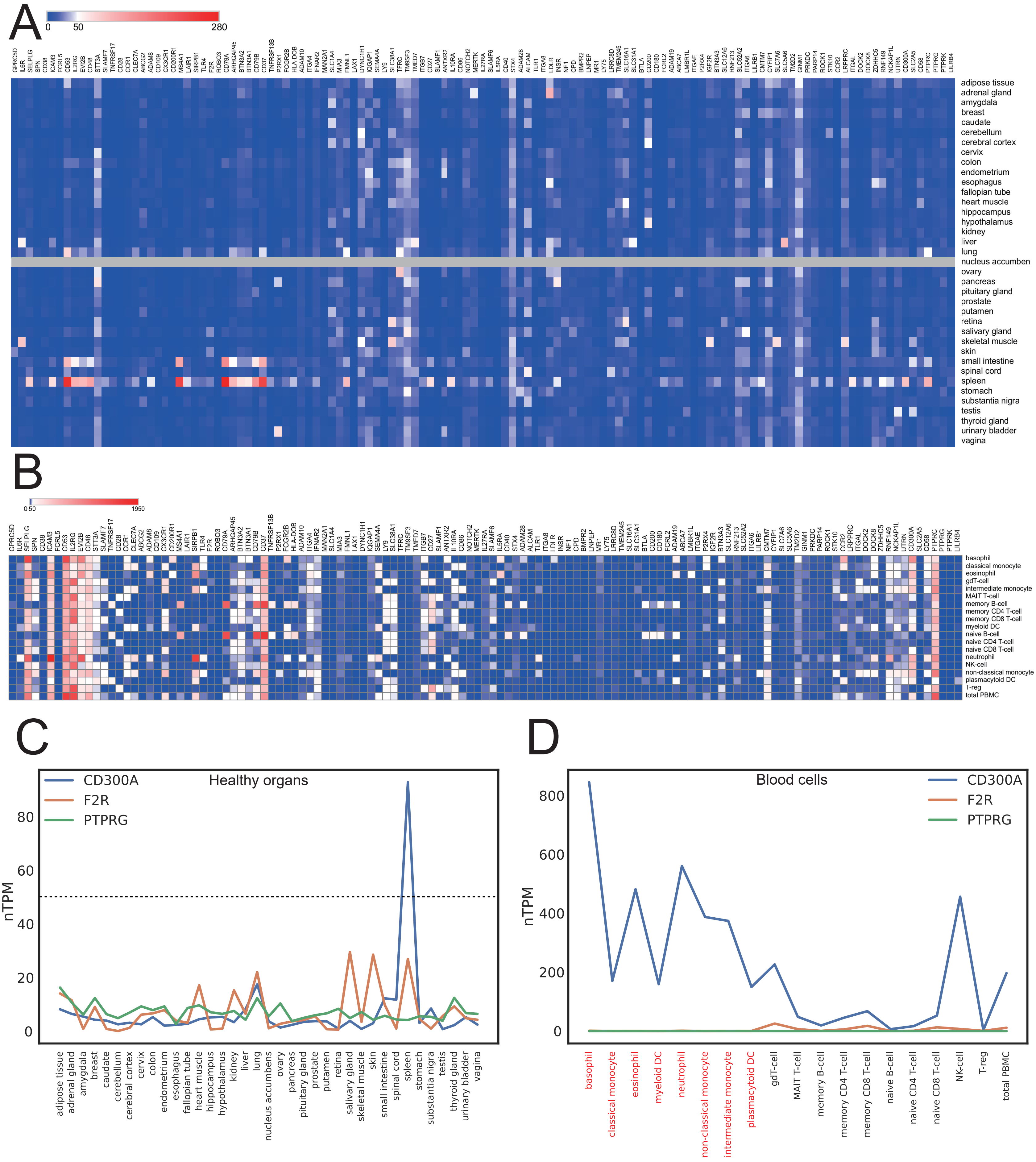
A
0
50
280
adipose tissue
adrenal gland
amygdala
breast
caudate
cerebellum
cerebral cortex
cervix
colon
endometrium
esophagus
fallopian tube
heart muscle
hippocampus
hypothalamus
kidney
liver
lung
nucleus accumben
ovary
pancreas
pituitary gland
prostate
putamen
retina
salivary gland
skeletal muscle
skin
small intestine
spinal cord
spleen
stomach
substantia nigra
testis
thyroid gland
urinary bladder
vagina
B
0 50
1950
basophil
classical monocyte
eosinophil
gdT-cell
intermediate monocyte
MAIT T-cell
memory B-cell
memory CD4 T-cell
memory CD8 T-cell
myeloid DC
naive B-cell
naive CD4 T-cell
naive CD8 T-cell
neutrophil
NK-cell
non-classical monocyte
plasmacytoid DC
T-reg
total PBMC
C
CD300A
F2R
PTPRG
Healthy organs
nTPM
0
20
40
60
80
adipose tissue
adrenal gland
amygdala
breast
caudate
cerebellum
cerebral cortex
cervix
colon
endometrium
esophagus
fallopian tube
heart muscle
hippocampus
hypothalamus
kidney
liver
lung
nucleus accumbens
ovary
pancreas
pituitary gland
prostate
putamen
retina
salivary gland
skeletal muscle
skin
small intestine
spinal cord
spleen
stomach
substantia nigra
testis
thyroid gland
urinary bladder
vagina
D
Blood cells
CD300A
F2R
PTPRG
nTPM
0
200
400
600
800
basophil
classical monocyte
eosinophil
myeloid DC
neutrophil
non-classical monocyte
intermediate monocyte
plasmacytoid DC
gdT-cell
MAIT T-cell
memory B-cell
memory CD4 T-cell
memory CD8 T-cell
naive B-cell
naive CD4 T-cell
naive CD8 T-cell
NK-cell
T-reg
total PBMC

**Supplementary Figure 6. Characteristics of candidates identified from high-risk subtypes.** **A**. Expression level of 125 candidates in healthy organs documented in the human protein atlas (THPA). **B**. Expression level of 125 candidates in blood cells documented in THPA. **C**. Three candidates identified in PR subgroup and their expression in healthy organs. **D**. Three candidates identified in PR subgroup and their expression in blood cells.

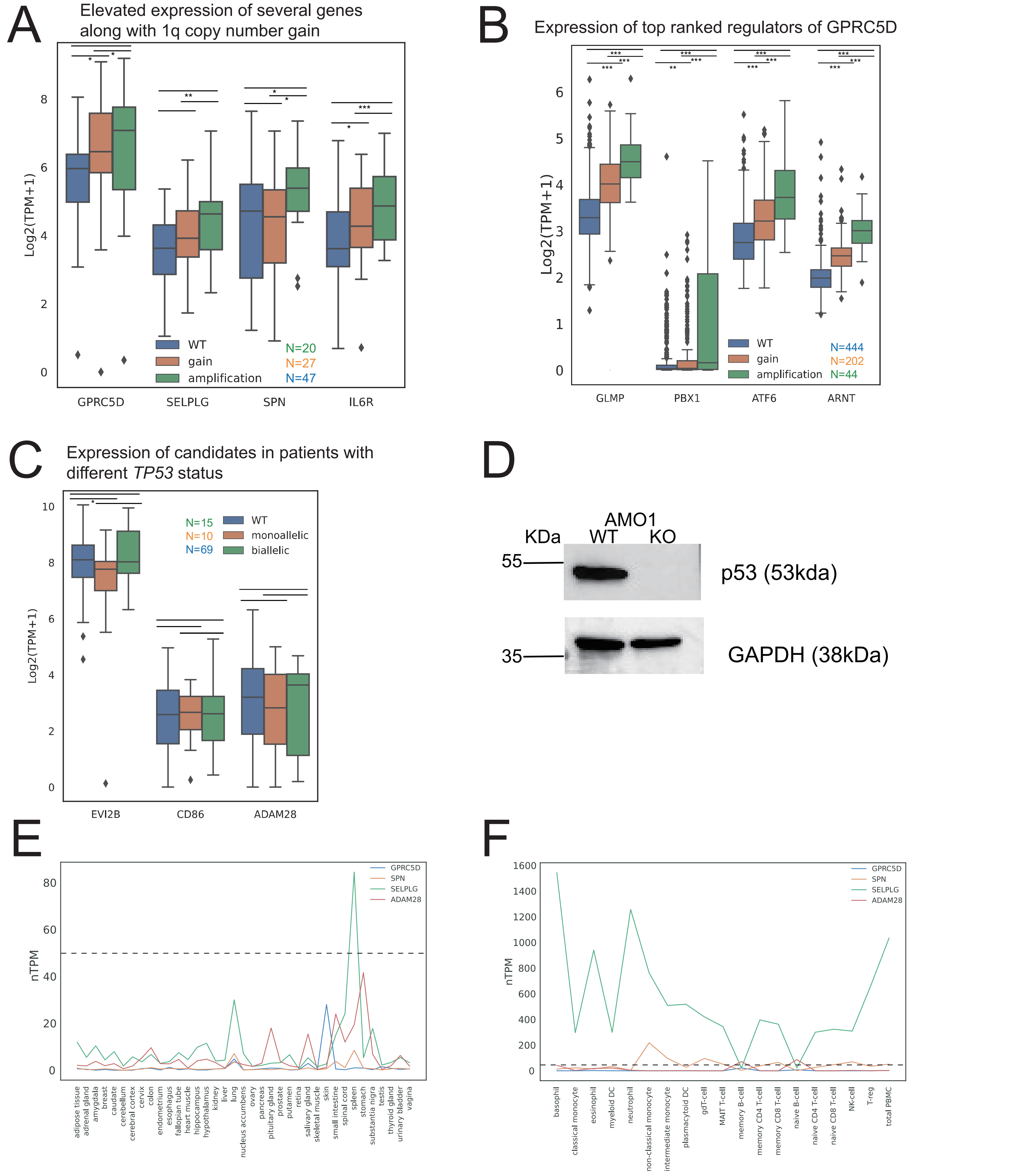
A
Elevated expression of several genes along with 1q copy number gain
Log2(TPM+1)
WT
gain
amplification
N=20
N=27
N=47
GPRC5D
SELPLG
SPN
IL6R
B
Expression of top ranked regulators of GPRC5D
Log2(TPM+1)
WT
gain
amplification
N=444
N=202
N=44
GLMP
PBX1
ATF6
ARNT
C
Expression of candidates in patients with different TP53 status
Log2(TPM+1)
N=15
N=10
N=69
WT
monoallelic
biallelic
EVI2B
CD86
ADAM28
D
AMO1
KDa
WT
KO
55
35
p53 (53kda)
GAPDH (38kDa)
E
nTPM
GPRC5D
SPN
SELPLG
ADAM28
adipose tissue
adrenal gland
amygdala
breast
caudate
cerebellum
cerebral cortex
cervix
colon
endometrium
esophagus
fallopian tube
heart muscle
hippocampus
hypothalamus
kidney
liver
lung
nucleus accumbens
ovary
pancreas
pituitary gland
prostate
putamen
retina
salivary gland
skeletal muscle
skin
small intestine
spinal cord
spleen
stomach
substantia nigra
testis
thyroid gland
urinary bladder
vagina
F
nTPM
GPRC5D
SPN
SELPLG
ADAM28
basophil
classical monocyte
eosinophil
myeloid DC
neutrophil
non-classical monocyte
intermediate monocyte
plasmacytoid DC
gdT-cell
MAIT T-cell
memory B-cell
memory CD4 T-cell
memory CD8 T-cell
naive B-cell
naive CD4 T-cell
naive CD8 T-cell
NK-cell
T-reg
total PBMC

**Supplementary Figure 7. Expression of selected candidates and their potential regulators**

**A**. Elevated expression level of four candidates as 1q copy number gain in IU cohort. **B**. Elevated expression level of potential regulators of *GPRC5D* on chromosome 1q. **C**. Expression level of biallelic *TP53*-specific candidates in IU cohort. **D**. A western blot result indicating the p53 protein expression between a *TP53* knockout and a WT cell-line of AMO1. **E.** Expression of selected candidates in healthy organs. **F.** Expression of selected candidates in blood cells. Statistical test in **A**, **B** and **C**: two-sided Mann Whitney U test; significance level: * $p<0.05$, ** $p<0.01$, *** $p<0.001$.

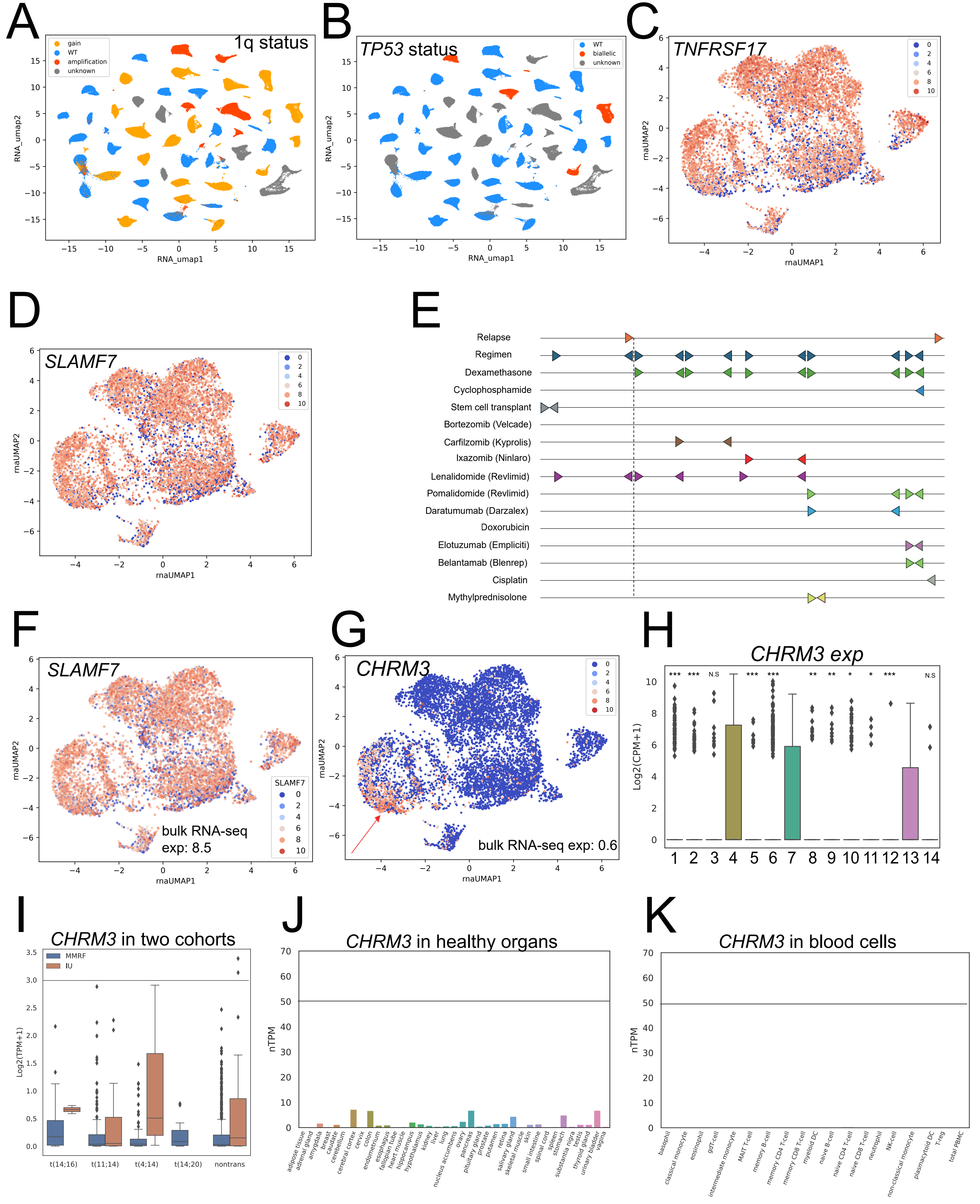
A
1q status
gain
WT
amplification
unknown
RNA_umap1
RNA_umap2
B
TP53 status
WT
biallelic
unknown
RNA_umap1
RNA_umap2
C
TNFRSF17
rnaUMAP1
rnaUMAP2
D
SLAMF7
rnaUMAP1
rnaUMAP2
E
Relapse
Regimen
Dexamethasone
Cyclophosphamide
Stem cell transplant
Bortezomib (Velcade)
Carfilzomib (Kyprolis)
Ixazomib (Ninlaro)
Lenalidomide (Revlimid)
Pomalidomide (Revlimid)
Daratumumab (Darzalex)
Doxorubicin
Elotuzumab (Empliciti)
Belantamab (Blenrep)
Cisplatin
Mythylprednisolone
F
SLAMF7
bulk RNA-seq
exp: 8.5
rnaUMAP1
rnaUMAP2
G
CHRM3
bulk RNA-seq exp: 0.6
rnaUMAP1
rnaUMAP2
H
CHRM3 exp
Log2(CPM+1)
*** *** N.S *** *** ** ** * * *** N.S
1 2 3 4 5 6 7 8 9 10 11 12 13 14
I
CHRM3 in two cohorts
MMRF
IU
Log2(TPM+1)
t(14;16)
t(11;14)
t(4;14)
t(14;20)
nontrans
J
CHRM3 in healthy organs
nTPM
adipose tissue
adrenal gland
amygdala
breast
caudate
cerebellum
cerebral cortex
cervix
colon
endometrium
esophagus
fallopian tube
heart muscle
hippocampus
hypothalamus
kidney
liver
lung
nucleus accumbens
ovary
pancreas
pituitary gland
prostate
putamen
retina
salivary gland
skeletal muscle
skin
small intestine
spinal cord
spleen
stomach
substantia nigra
testis
thyroid gland
urinary bladder
vagina
K
CHRM3 in blood cells
nTPM
basophil
classical monocyte
eosinophil
gdT-cell
intermediate monocyte
MAIT T-cell
memory B-cell
memory CD4 T-cell
memory CD8 T-cell
myeloid DC
naive B-cell
naive CD4 T-cell
naive CD8 T-cell
neutrophil
NK-cell
non-classical monocyte
plasmacytoid DC
T-reg
total PBMC

**Supplementary Figure 8. Characteristics of candidates' expression in subclones.**

**A**. 1q status of 49 patients with single-cell RNA-seq. **B**. *TP53* status of 49 patients with single-cell RNA-seq. **C**. single-cell expression of *TNRFSRF17*/BCMA among subclones of 49 patients. **D**. single-cell expression of *SLAMF7*/CD319 among subclones. **E**. Treatment timeline of a relapsed t(4;14) patient. **F**. expression level of *SLAMF7*/CD319 in the relapsed t(4;14) patient. **G**. *CHRM3* expression among subclones of this patient. **H**. Differential expression of *CHRM3* among subclones of this patient. **I**. Expression level of *CHRM3* among primary subtypes in bulk RNA-seq data. **J**. expression level of *CHRM3* in healthy organs. **K**. expression level of *CHRM3* in blood cells. Statistical test in **H**: Mann Whitney U test. Significance level: * $p<0.05$, ** $p<0.01$, *** $p<0.001$.

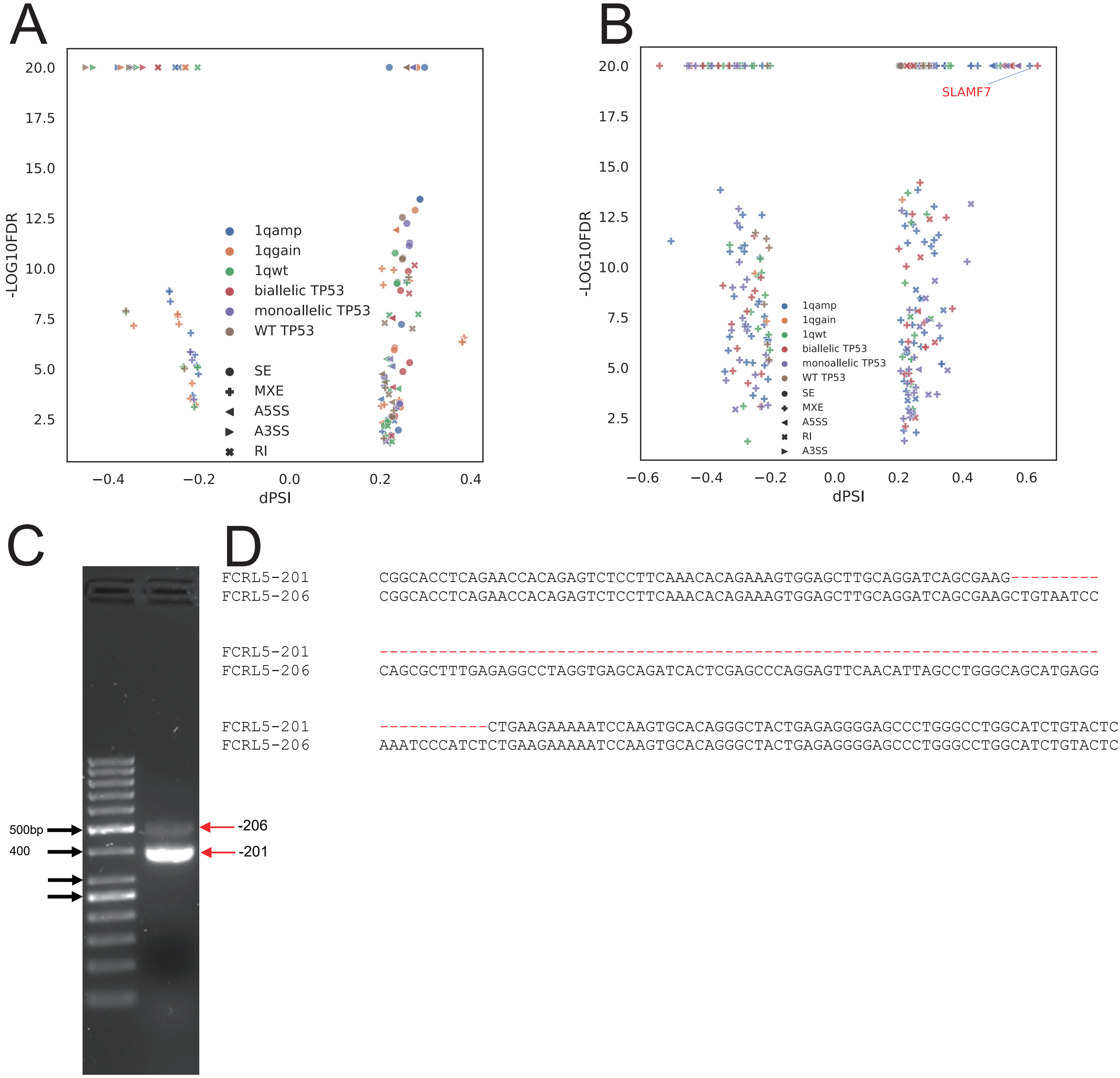
A
-LOG10FDR
dPSI
1qamp
1qgain
1qwt
biallelic TP53
monoallelic TP53
WT TP53
SE
MXE
A5SS
A3SS
RI
B
-LOG10FDR
dPSI
SLAMF7
1qamp
1qgain
1qwt
biallelic TP53
monoallelic TP53
WT TP53
SE
MXE
A5SS
RI
A3SS
C
500bp
400
-206
-201
D
FCRL5-201 CGGCACCTCAGAACCACAGAGTCTCCTTCAAACACAGAAAGTGGAGCTTGCAGGATCAGCGAAG---------
FCRL5-206 CGGCACCTCAGAACCACAGAGTCTCCTTCAAACACAGAAAGTGGAGCTTGCAGGATCAGCGAAGCTGTAATCC
FCRL5-201 -------------------------------------------------------------------------
FCRL5-206 CAGCGCTTTGAGAGGCCTAGGTGAGCAGATCACTCGAGCCCAGGAGTTCAACATTAGCCTGGGCAGCATGAGG
FCRL5-201 -----------CTGAAGAAAAATCCAAGTGCACAGGGCTACTGAGAGGGGAGCCCTGGGCCTGGCATCTGTACTC
FCRL5-206 AAATCCCATCTCTGAAGAAAAATCCAAGTGCACAGGGCTACTGAGAGGGGAGCCCTGGGCCTGGCATCTGTACTC

**Supplementary Figure 9. Alternative splicing landscapes of 125 high-risk identified candidates.**

**A**. Alternative splicing landscapes among 125 high-risk candidates in MMRF cohort. **B**. Alternative splicing landscapes among 125 high-risk candidates in IU cohort. **C**. A RT-PCR experiment result indicating the expression of *FCRL5*-206. Columns in (**C**): 1. A PDX sample; 2. A SACHI cell-line 3. No reverse transcriptase control; 4. NTC (PCR negative control). Rows in (**C**): fragment size. Expected fragment size: cryptic exon: 495 bps; WT: 404 bps. **D**. cDNA sequences of exon 4, the cryptic exon (dashed basepairs) and exon3.

**Supplementary Table legends**

**Supplementary Table 1. Summary of candidates identified under all conditions.**

**Supplementary Table 2. Association between expression of candidate found in population and patient survival.**

**Supplementary Table 3. Association between expression of candidate found in primary subtypes and patient survival.**

**Supplementary Table 4. Association between expression of candidate found in high-risk subtypes and patient survival.**

**Supplementary Table 5. Primer designing in TP53 knockout experiments.**
